# Emergence and spread of feline infectious peritonitis due to a highly pathogenic canine/feline recombinant coronavirus

**DOI:** 10.1101/2023.11.08.566182

**Authors:** Charalampos Attipa, Amanda S Warr, Demetris Epaminondas, Marie O’Shea, Andrew J Hanton, Sarah Fletcher, Alexandra Malbon, Maria Lyraki, Rachael Hammond, Alexandros Hardas, Antria Zanti, Stavroula Loukaidou, Michaela Gentil, Danielle Gunn-Moore, Stella Mazeri, Christine Tait-Burkard

**Author notes:** Corresponding authors: (Epidemiology, pathology), (Sequencing, bioinformatics), (General enquiries, sequence analysis, virology, structure). Contributed equally to this work. CA was responsible for epidemiology, pathology and sample/outbreak data collection. AW performed sequencing and data analysis thereof. Both are equally important, but the latter couldn’t have happened without the first.

## Abstract

Cross-species transmission of coronaviruses (CoVs) poses a serious threat to both animal and human health^1–3^. Whilst the large RNA genome of CoVs shows relatively low mutation rates, recombination within genera is frequently observed and demonstrated^4–7^. Companion animals are often overlooked in the transmission cycle of viral diseases; however, the close relationship of feline (FCoV) and canine CoV (CCoV) to human hCoV-229E^5,8^, as well as their susceptibility to SARS-CoV-2^9^ highlight their importance in potential transmission cycles. Whilst recombination between CCoV and FCoV of a large fragment spanning orf1b to M has been previously described^5,10^, here we report the emergence of a novel, highly pathogenic FCoV-CCoV recombinant responsible for a rapidly spreading outbreak of feline infectious peritonitis (FIP), originating in Cyprus^11^. The minor recombinant region, spanning spike (S), shows 96.5% sequence identity to the pantropic canine coronavirus NA/09. Infection has rapidly spread, infecting cats of all ages. Development of FIP appears very frequent and sequence identities of samples from cats in different districts of the island is strongly supportive of direct transmission. A near cat-specific deletion in the domain 0 of S is present in >90% of FIP cats. It is unclear as yet whether this deletion is directly associated with disease development and may be linked to a biotype switch^12^. The domain 0 deletion and several amino acid changes in S, particularly the receptor binding domain, indicate potential changes to receptor binding and cell tropism.

## Main

Following two epidemics, SARS-CoV (2002-4) and MERS-CoV (2012-ongoing), and a pandemic of previously unseen proportions, SARS-CoV-2 (2019 onwards), coronaviruses no longer need lengthy introductions of importance and scale. They are not only present in the human population but also in wildlife^13–15^, companion animals^16–19^ and livestock^13,20,21^, and in all species these viruses have major impacts. The innate ability of coronaviruses to recombine with other coronaviruses continues to fuel their species-switching ability. It is therefore not surprising that both human and animal coronaviruses are linked in complex transmission and evolution cycles^3,14,19,22^.

FCoV is found across the globe. The virus exists in two biotypes with the main biotype, feline enteric coronavirus (FECV), showing low virulence, and clinical signs are typically limited to mild enteritis. The second biotype of FCoV, proposed to originate each time from a mutation in an FECV-infected cat (reviewed in ^23^), is known as feline infectious peritonitis virus (FIPV). FIPV causes feline infectious peritonitis (FIP), which is a fatal disease if left untreated. Clinical signs include an abdomen swollen due to peritoneal fluid, fever, weight loss, lethargy, anorexia, dyspnea, ocular abnormalities and neurological signs^8,16,24,25^. Mutations in S or the accessory genes 3abc and 7ab of FCoV^8,16,23,26^ are thought to result in changes to the virus’s tropism from cells in the enteric tract to macrophages, resulting in the different disease presentation seen with the two biotypes. This change in primary tropism also impacts the virus’s ability to transmit from cat to cat, with the main transmission pathway of FECV being fecal-oral and FIPV typically having relatively poor transmission potential. Antivirals, including remdesivir and GS-441524 have recently been successfully used to treat cats with FIP^27^.

FCoV and CCoV both belong to the *Alphacoronavirus 1* species alongside the porcine transmissible gastroenteritis virus (TGEV) and, probably, the never fully sequenced rabbit enteric coronavirus^28,29^. Both FCoV and CCoV have evolved two different serotypes through complex recombination events between the two viruses with a suspected gradual evolution from CCoV-II to TGEV and the later S deletion to porcine respiratory coronavirus (PRCV)^23,30^. Whilst recombination events between CCoV and FCoV have significantly contributed to the serotype evolution and have been frequently described, so far, none of them led to enhanced transmissibility of FIP beyond closest contact^10,23,31,32^. Similarly, recombination events have been reported between CCoV and TGEV^33^, the latter has been found to recombine with the pedacoronavirus (alphacoronavirus genus) porcine epidemic diarrhea virus^6^. These observations are particularly important in view of the *Alphacoronavirus 1*-related human infections recently observed^19,22^.

In 2023, we alerted the veterinary field to an outbreak of FIP in Cyprus, where there had been a concerning increase in cases^11^. Cases were recorded as FIP only if they had compatible clinicopathological signs and a positive RT-qPCR for FCoV in one of the following samples; peritoneal fluid, pleural fluid, cerebrospinal fluid, fine needle aspiration biopsies, or tissue biopsies from granulomatous lesions or positive FCoV immunohistochemistry (IHC) of granulomatous lesions (IHC). RT-qPCR confirmed FIP cases had increased from three and four in 2021 and 2022, respectively, to 215 cases (186 thereof in 2023) from January 2023 to June 2024, representing more than a 50-fold increase. The outbreak emerged in January 2023 in Nicosia, the capital of Cyprus. By February the area of Nicosia recorded the peak number of cases in any district (Figure 1A & B). The next highest number of cases was observed in Famagusta, which peaked in March, and by then the outbreak had spread to all districts of the Republic of Cyprus. (Supplementary Tables S1-6). In June and July there was a decline in RT-qPCR confirmed cases, which coincided with a large media awareness campaign to alert veterinarians to the spread of FIP^11^. This fall in cases was likely due to most veterinarians diagnosing cases based on clinicopathological findings without performing PCR testing due to the additional financial cost. On August 3^rd^, the Republic of Cyprus minister’s cabinet approved the use of human coronavirus medication stocks in cats with FIP. For veterinarians to have access to this medication, amongst others, a PCR confirmation was required, reflecting the increased cases recorded during August 2023 (Figure 1A & B). The number of unreported cases in Cyprus remains very high, not least due to the high number of feral cats. Estimates from the Pancyprian Veterinary Association indicate approximately 10,000 cases were presented in the veterinary clinics from January 2023 until July 2023 due to clinical signs of FIP. Furthermore, in October 2023, a first imported case of FIP was confirmed in the UK^34^. The most common clinical form of FIP was effusive (63.72%; Figure 1C), followed by neurological FIP (26.05%) and the non-effusive (dry) form (10.23%) (Supplementary table S3). Whilst the initial wave of infections appeared to slow down in August 2023, a second wave can be observed September 2023-March 2024 with the beginning of a third wave indicated from June 2024. The only distinctive feature of the second and third waves is a higher proportion of non-effusive forms (Supplementary tables S3-S5). Possible explanations for this include: an altered immune response from the host following initial seroconversion and re-infection; that the non-effusive form takes longer to develop; or that cats presenting with the non-effusive form in the early stages of the outbreaks were not diagnosed due to their less common clinical signs.

**Figure 1:**
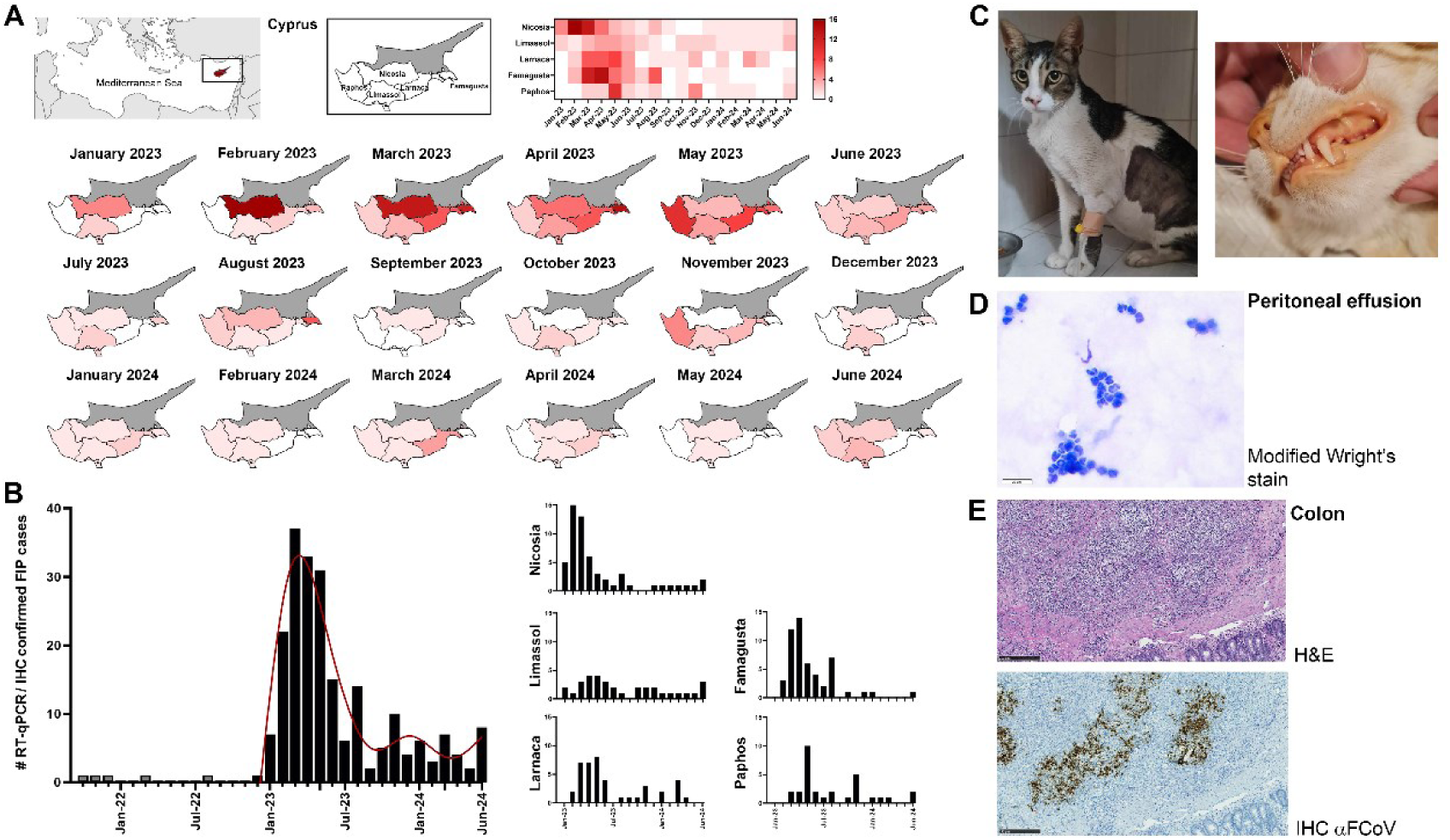
Epidemiology and Pathology of the FIP outbreak in Cyprus, 2023-June 2024. **A)** Distribution of RT-qPCR/IHCconfirmed FIP cases across Cyprus. First image locates Cyprus within the Eastern Mediterranean. Darker colors indicate higher numbers of confirmed cases over time within each district as highlighted in the overview heatmap with key. **B)** RT-qPCR/IHCconfirmed case rates resolved by time and province. A 6-knot spline interpolation highlights the three waves observed thus far. **C)** Clinical presentation of cats with FIP due to FCoV-23. Left; cat with the effusive form of FIP showing abdominal distention due to peritoneal effusion, an unkept coat, low body condition score, and poor muscle condition. Right; cat presenting with jaundice evidenced by yellow/orange discoloration of the mucous membranes and mucocutaneous junctions. Images courtesy of Dr Eva Georgiadi. **D)** Photomicrograph of fluid smear from a peritoneal effusion from a cat with confirmed FIP due to FCoV-23 infection. Non-degenerative neutrophils are present on a protein rich background shown using a Modified Wright’s stain. Scale bar represents 20µm. E) Photomicrographs showing a section of colonic mucosa and submucosa from a cat with confirmed FIP due to FCoV-23 infection. Top; H&E-stained histology section shows coalescing infiltration of predominantly the submucosa by aggregates of primarily neutrophils and macrophages surrounded by fewer lymphocytes and plasma cells. The muscularis mucosae is disrupted by the inflammation. Bottom; Immunohistochemistry staining against FCoV in a histology section mirroring the above. Extensive positive FCoV cytoplasmic staining for cells at the center of each aggregate/pyogranuloma in cells with macrophage-like morphology. Scale bar represents 200µm.

Where peritoneal or pleural fluid was assessed by cytology, non-degenerative neutrophils admixed with macrophages and small lymphocytes were seen in a protein rich background (Figure 1D). 17 cases were assessed by histopathology, including intestinal mass (n=8), lymph node (n=5), and kidney (n=4). All showed similar histological features, with multifocal to coalescing, pyogranulomatous to necrotizing and lymphoplasmacytic inflammation (Figure 1E). The angiocentric nature can be seen in some areas, whilst in others there is total effacement of the tissue. Immunohistology for FCoV antigen demonstrated a heavy viral load within intralesional macrophages (Figure 1E).

RNA samples were obtained from 163 confirmed FIP cases between 2021 and November 2023, representing a mixture of geographic origin, sex, and clinical presentation (Supplementary Tables S7-S12). There are inherent problems with obtaining sufficient read-depth with Illumina sequencing on FIPV RNA samples. Therefore, we opted for cDNA/PCR-amplification-based Nanopore sequencing to better understand the Cypriot outbreak and to determine if cat-to-cat transmission is occurring. An initial crude approach based on alignments of reposited FCoV-1 and FCoV-2 sequences yielded a mixed consensus sequence for the Cypriot outbreak strain, FCoV-23. The consensus was used to design a new primer scheme for targeted amplification of FCoV-23 for Oxford Nanopore-based sequencing. This yielded a total of 19 full-length genomes, 45 partial sequences spanning a section of orf1b (∼1,000 bp), 63 spanning the first part of S (∼2,250 bp), and 42 spanning ORFs 3c/E(4)/M(5) (∼1,000 bp). These include samples from two cats presenting with FIP following recent import from Cyprus to the UK. Other samples were degraded or contained too few viral copies. None of the seven samples from before 2023 amplified (Supplementary Tables S13-S18).

The partial S region sequence of the Cypriot/UK FCoV samples produced two distinct versions of the S sequence. BLAST was used to identify close relatives of these S sequences. The first S sequence is most closely related to an FCoV-1 (MT444152) with 79% similarity, which occurred in one Cypriot and one UK-import sample. The others are almost entirely CCoV-2 S, flanked by FCoV-1 sequence. The CCoV-2 sequence is most closely related to the NA/09 strain (JF682842), a hypervirulent pantropic canine coronavirus (pCCoV)^35^, at 96.5% sequence identity. As shown by maximum likelihood analysis the S sequence is also closely related to other pCCoV S sequences with only partial S sequences available (Figure 2, Supplementary Figure S1-S2). Interestingly, the majority of samples contained a deletion at the N-terminus of S, which is analyzed in more detail below. Except for the two samples highlighted above, all samples, including one UK-import case, align most closely with a pCCoV S. This is likely a defining feature of the virus circulating in the outbreak in Cyprus. In contrast, maximum likelihood trees of regions of ORF1b and ORF3c/E/M (Figure 2, Supplementary Figures S3-S4) show clustering of all sequences with FCoV-1. Whilst there appears to be more diversity in ORF1b, one must also consider the scale of the different trees.

**Figure 2:**
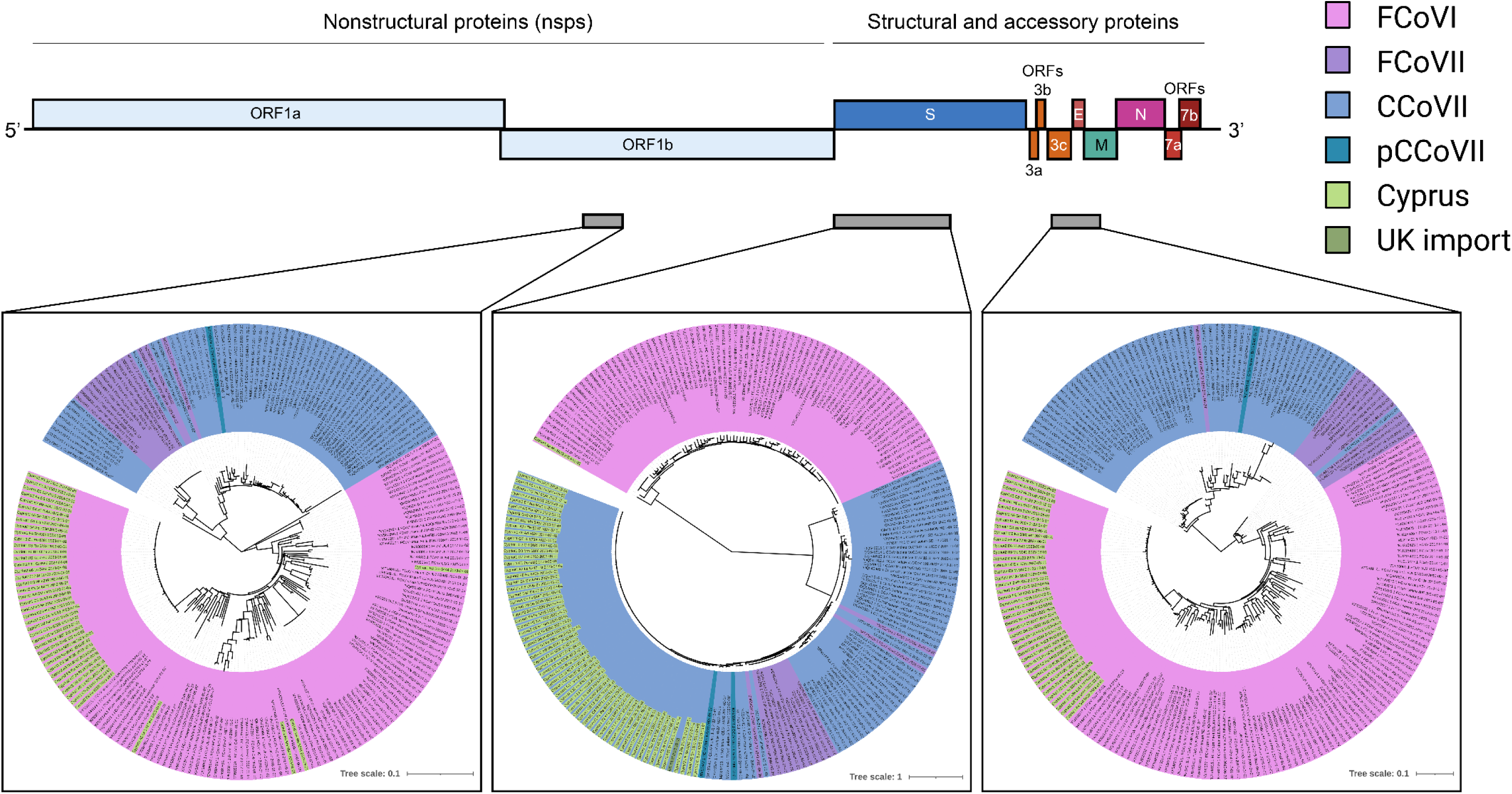
Genomic sequence analysis of Cypriot/UK-import FCoV cases. Sequences from three different genomic regions: Orf1b, S, and Orf3c/E/M, as indicated by the regions marked on the overview of the genome, were obtained through Nanopore sequencing from 45, 63, and 42, Cypriot and UK-import samples, respectively. Following initial BLAST analysis, maximum likelihood trees were generated including other FCoV and CCoV strains to assess genetic similarity of each region. CCoV-2 genomes are highlighted in blue with pCCoV-2 genomes within this region displayed in a darker blue. FCoV-1 genomes are highlighted in pink and FCoV-2 genomes in purple. Samples from Cyprus (green) and one UK-imported Cypriot cat (dark green) can be seen clustered with FCoV-1 sequences in the Orf1b and Orf3c/E/M regions. The S gene, however, clusters with CCoV-2, most closely with pCCoV-2. Different numbers of sequences are present in each tree due to missing sequence or poor sequence quality and/or alignments in genomes downloaded from NCBI, and due to not all regions being sequenced in all individuals from our samples.

A recombination analysis was carried out comparing the FCoV-23 genome using representative full-length sequence 2-C11 Re 10276 with FCoV-1, FCoV-2 and CCoV-2 strain CB/05, since NA/09 is not available as a complete genome. Figure 3A visualizes the Bootscan^36^ analysis, the RDP5^37^ analysis and the pairwise distances between the sequences. Significant p-values supporting the recombination were reported by multiple methods as listed in Supplementary Table S19. The MaxChi^38^ breakpoint matrix is available in Supplementary Figure S6. The recombination is very clear and includes a small region of the ORF1b gene and the majority of S with breakpoints around location 18,420 and 23,880. Figure 3B shows the historical break points between FCoV-1 and CCoV-2 that created FCoV-2 alongside the recombination identified in FCoV-23.

**Figure 3:**
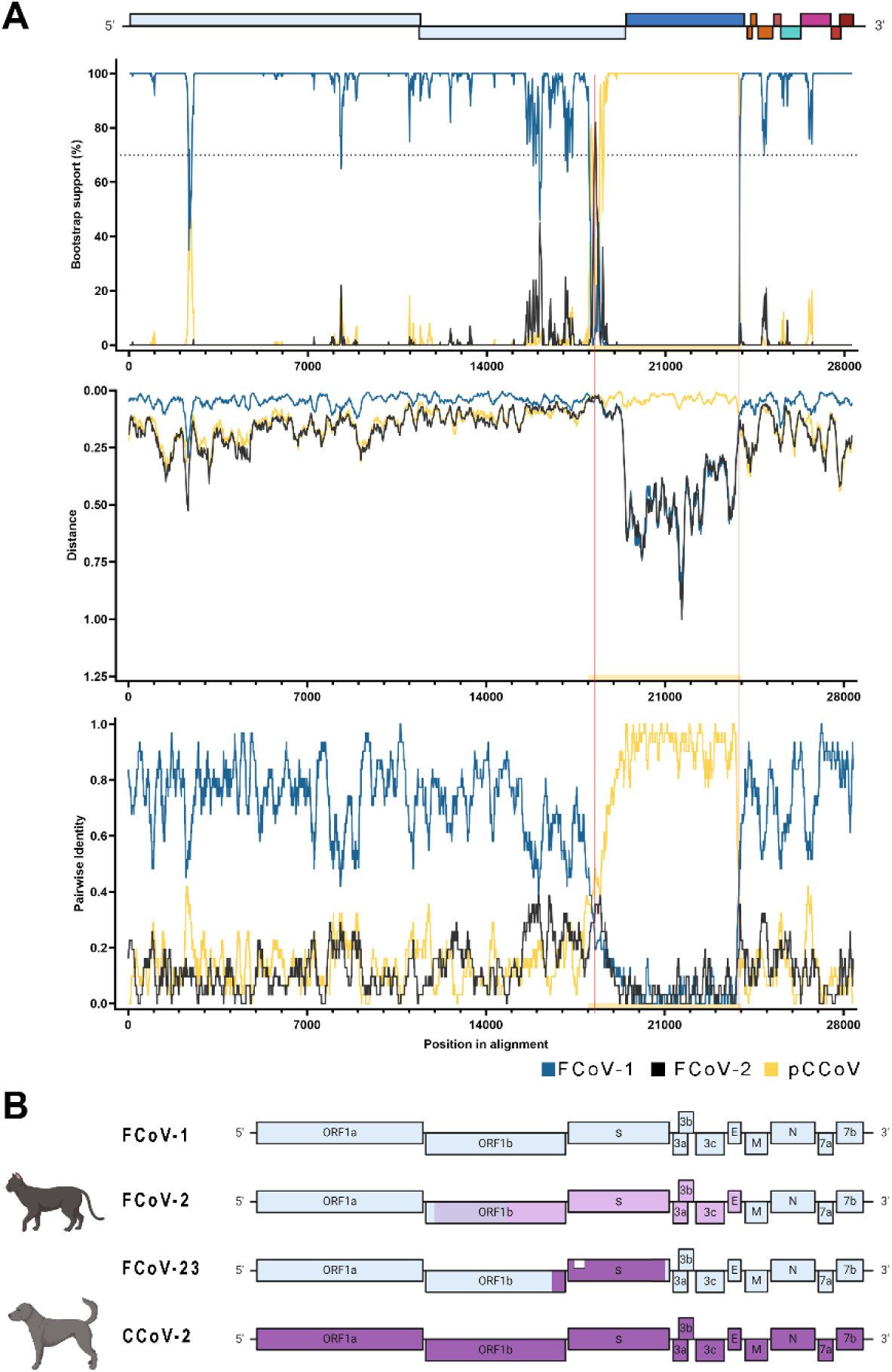
Recombination analysis. **A)** Visualizations of a recombination analysis carried out on the assembled FCoV-23 genome 2-C11 Re 10276 and representative genomes of FCoV-1 (blue), FCoV-2 (black) and pCCoV (yellow). The yellow panel shows the likely recombination break region, with a red vertical line showing the likely break point. The first panel shows the results of the Bootscan analysis, the second panel shows the sequence distance, and the third panel shows the RDP pairwise identity analysis. All three panels show good support for the recombination between FCoV-1 and pCCoV. These results are further supported by high statistical likelihood of recombination shown in Supplementary Table S21. Also see Supplementary Figures S6&7 and Table S21 for recombination analysis of a representative domain 0-deletion FCoV-23. **B)** Schematic representation of major recombinations observed in FCoV and CCoV. A common ancestral origin virus is thought to have given rise to the original FCoV-1 and CCoV-1 types with further evolution in CCoVs yielding CCoV-2 and CCoV-1/2 serotypes. FCoVs are thought to have recombined with CCoV-2 to form FCoV-2. FCoV-2/CCoV-2 recombinations have shown to have different recombination points as highlighted by the different colorations (reviewed in 23). The Cypriot FCoV strain, termed FCoV-23, is a recombinant between an FCoV-1 strain and a S recombination with a pantropic CCoV-2, pCCoV. Furthermore, deletion variants are observed in a majority of sequenced cases (white box and Figure 4).

#Supplementary Figure S5 furthermore shows a neighbor joining tree for FCoV-23 along with other members of *Alphacoronavirus 1* and a distantly related Canine Respiratory Coronavirus as an outgroup. The assembled genome clusters with representatives of FCoV-1, similar to the clustering of the amplicons outside of S.

The main determinant in disease development and transmission of FCoV-23 appears to be the S recombination. One of the main suggested determinants of biotype changes, the furin cleavage site (FCS) at the S1/S2 interface^23,26^ is absent in FCoV-2 and also FCoV-23. An interesting observation, however, is that the majority of samples (>90%) show a deletion in domain 0 (ΔD0), strongly resembling the deletion observed in TGEV and PRCV (Figure 4A). Interestingly, whilst the rest of the genome of FCoV-23 is highly conserved, the domain 0 deletion appears to be almost cat-specific (Figure 4B). This indicates that this mutation occurs in host rather than being transmitted. However, the only two fecal samples (also affected by sampling bias; <6% of our samples are feces-derived) where sequences could be obtained show ΔD0. An in-host mutation further begs the question of whether ΔD0 is associated with disease development. Unfortunately, all our samples so far are from symptomatic, confirmed FIPV cases and further investigation is required to answer this question.

**Figure 4:**
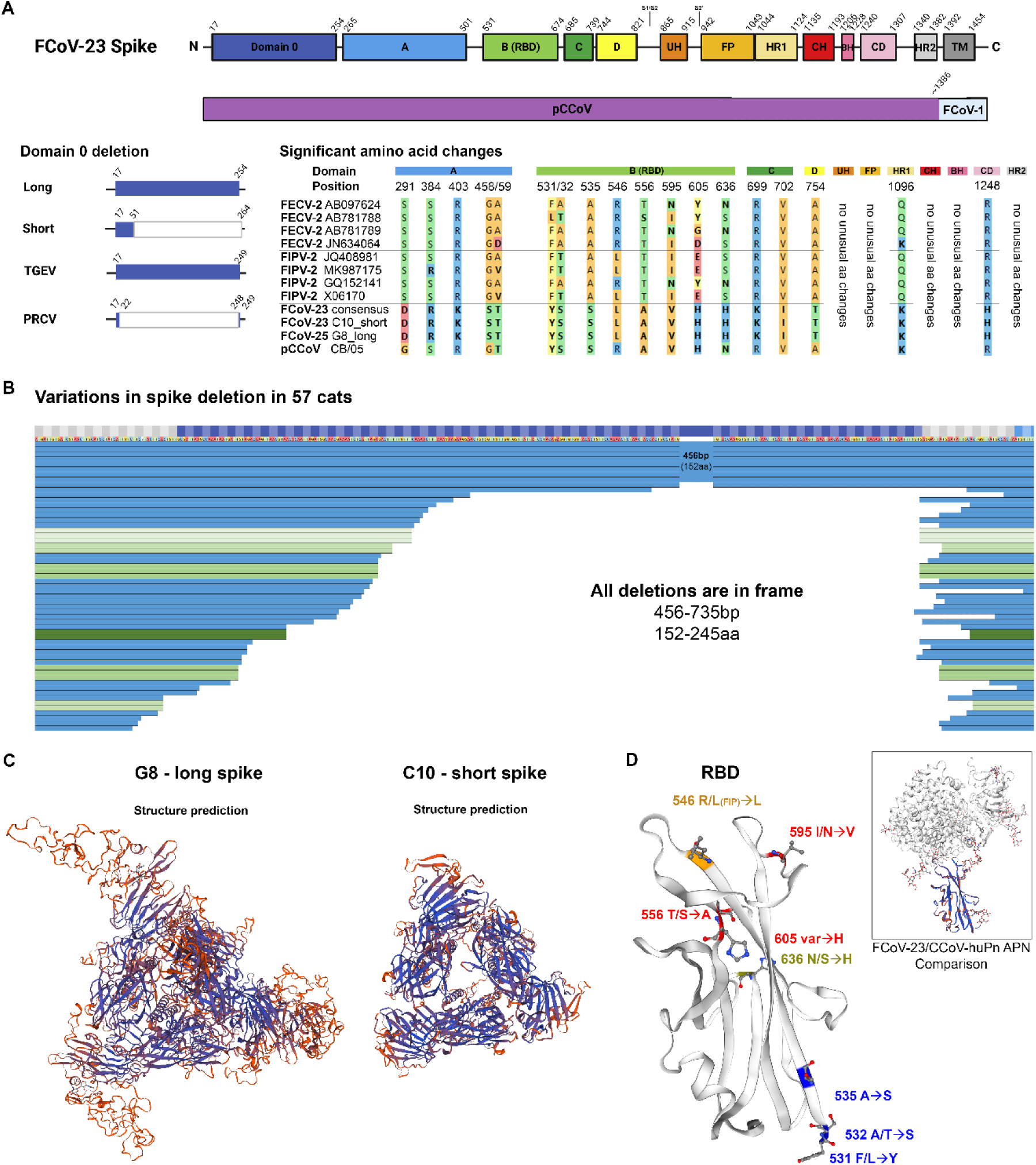
Protein sequence and structural analysis of FCoV-23 Spike. **A)** Analysis of the different domains of FCoV-23 S. Domain 0 in its full-length version shows high similarity to CB/05 pCCoV; however most prominent is the large, variable deletion between aa11-265 strongly resembling the deletion previously observed between the TGEV and PRCV S. Multi-sequence alignment was performed against four confirmed FECV-2 and four confirmed FIPV-2 mirroring sequences used for computational analysis by Zehr *et al.*^26^. Highlighted are amino acid changes with significant change against FCoV-2s and also position 546, where a Leucin is more predominant in FIPV rather than FECV sequences. Colors are following the RasMol amino acid color scheme. **B)** Domain 0 S deletions from 57 cats (8 full length) are shown from the beginning of S to amino acid 267. Sequences are ordered by the shortest 3’/N-terminal sequence. Identical deletions are highlighted in shades of green. All deletions show a multiplicity of 3 resulting in in-frame deletions. **C)** Structural modelling of the “full-length” or long S version, represented by sample G8, and the “domain 0 deletion” or short version, represented by sample C10. Samples were modelled against the CCoV-HuPN-2018, experimentally determined S in the swung out confirmation 7usa.1.A^40^. The proximal confirmation and comparisons may be found in Supplementary Figure 8. Colors indicate confidence with blue highlighting strong and red highlighting weak confidence. **D)** Modelling or amino acid changes on a structural prediction of the FCoV-23 RBD against CCoV-HuPN-2018 7u8I.1.B. Significant amino acid changes as identified in A) are highlighted with side-chains in blue for variations to FCoVIIs distant from the RBD, red for variations close to the RBD, orange for the variation at aa546 similar to FIPV-2, and olive for the variation differing both from FCoVII and pCCoV. A comparison showing a confidence model of the FCoV-23 RBD structure prediction paired with the CCoV-HuPN-2018-canine aminopeptidase N (APN) is shown for orientation and binding visualization in the top right corner. Colors indicate confidence with blue highlighting strong and red highlighting weak confidence.

In other coronaviruses, including TGEV^39^ and CCoV-HuPn-2018^40^, domain 0 was shown to bind sialosides. Modelling the structure of S against the closely related, experimentally confirmed CCoV-HuPn-2018 S^40^ shows a much more compact conformation for ΔD0 S and similarity to a structural prediction based on a “swung-out” or a “proximal” confirmation template (Figure 4C, Supplementary Figure S8). A number of amino acid changes were observed between “classical” FECV-2 and FIPV-2 S. In particular domains A, B, and the receptor binding domain (RBD) show a number of class-changing amino acid changes distinct from FCoV-2 S (Figure 4A). Modelling the RBD changes against the structure highlights changes at positions 546 and 595, as well as 556, 603, and 636 as being potentially strongly influential to receptor binding properties (Figure 4D).

Previously indicated key proteins for biotype switch S, Orf3abc and 7b were compared to the recently published computational analysis of mutations observed in FECV versus FIPV by Zehr *et al.*^26^. The suggested key determinant of FIPV in the FCoV-2 S, position 1404, shows a new amino acid, leucine instead of valine, in FCoV-23. However, this section of spike is derived from FCoV-1. It is therefore unclear how this recombination may impact overall functionality in the context of the rest of S being FCoV-2-like. Other positions in S show a mixture of FIPV versus FECV tendencies (Supplementary Table S23). Whilst a new mutation was identified in Orf3a and b, no specific indications of pathogenesis could be determined (Supplementary Table S24). Similarly, in Orf7b, two new mutations but no indication of pathogenesis could be identified (Supplementary Table S25).

## Discussion

Recombination within the *Alphacoronavirus 1* species has been frequently observed previously and has even given rise to the FCoV-2 serotype^5,23,24^(Figure 3B) in cats. Even complex recombination between FCoV-1 and CCoV has previously been observed^10^. However, in this work, we have identified a new subtype of FCoV that has recombined with a hypervirulent strain of pCCoV. The recombinant, which we propose naming FCoV-23, shows clearly distinct properties from previously observed FCoV infections. The sequence similarity found between FCoV-23 and CCoVII is higher than would be expected without an additional, recent recombination event.

Our data suggest that there is direct transmission of FCoV-23 between cats based on high sequence identity (>99.17% genome-scale including varying S-deletion length, Supplementary figures S9 & S10), high viral loads (RT-qPCR only) in feces, as well as the wave-like movement of disease across the island. However, it is unclear whether this virus still needs a biotype change to result in FIP. A potential reason for differences in pathogenicity may be the acquiring of a deletion of S domain 0. This is observed in >90% of FIP cases studied here and in a near host-specific pattern. More work is needed to assess the properties of ΔD0 S, as well as investigation into asymptomatic carriers. Whilst alphacoronaviruses show great cross-neutralizing activity^40^, the onset of FIP appears rapid and shows little discrimination in age of the infected cats highlighting that FCoV-23 is able to circumnavigate preexisting immunity. If a biotype change is required, it clearly happens far more frequently in FCoV-23 than any of the previously observed FCoV infections. We therefore must study FCoV-23 to gain further understanding of the FECV/FIPV change. The scale of this FIP outbreak has not been observed previously. Concerningly, Cyprus has a high population of unowned cats that are frequently relocated to other parts of Europe, and the wider world. The risk of spreading this outbreak is significant as evidenced by the first confirmed UK case^34^.

CCoV infections in dogs are typically self-limiting, producing mild enteritis or presenting asymptomatically. Previous work on the FCoV-23 close relative strain, CCoV NA/09 and CB/05, found that the virus was hypervirulent and infected a range of organs in canine hosts. CCoV typically infects cells of the enteric tract, but pCCoVs were shown to spread to a range of internal organs, including intestine, lung, spleen, liver, kidney, lymph nodes, brain, and even T and B cells ^35,41–43^. CB/05 has also been identified as responsible for small outbreaks^41^.

The clinical signs of FCoV-23 are similar to those observed in classical FIP cases. However, FIP infection itself already shows a strong pantropism, affecting many organs in infected cats^25^. What is remarkable is the high number of FCoV-23 positive cells observed in immunohistochemistry samples (Figure 1C). This is indicative of high viral loads, but must be validated through quantification. Furthermore, a relatively high percent (Supplementary Table 4) of confirmed cases in Cyprus was presented with neurological signs (26%), which is almost double of what would be expected with classical FIP (14%)^27^. This may be due to increased awareness of presentation or be inherent to FCoV-23 neurotropism; however, further investigation is required.

Significant changes to the S protein in FCoV-23 may provide some clues as to the enhanced pathogenicity of this virus. FCoV-23, like other FCoV-2s and CCoV-2, does not contain an FCS at the S1/S2, and the related virus CCoV-HuPn-2018 was experimentally found to be uncleaved^40^. Uncleaved S has been shown to be more stable than cleaved and could enhance fecal shedding and stability in the environment^44^. Conversely, the cleavage at S1/S2 may facilitate the movement of the N-terminal domains and allow the RBD to adopt the receptor-binding-competent form. However, the deletion of domain 0 may compensate for the increased rigidity. Whilst a similar deletion between TGEV and PRCV was initially indicated as the determinant between a primarily enteric and primarily respiratory tropism, respectively^45,46^, a recent study shows a different result *in vivo*^47^. Sialoside binding is an important feature of coronavirus infections and may contribute to intracellular spread^48^. It is then all the more surprising that FCoV-23 loses the sialoside-binding domain 0. However, whilst binding to sialosides can enhance virus attachment and entry into host cells, binding to sugars can lead to increased retention of virus on producer cells. Whilst some viruses, like influenza, solve this issue by encoding their own sugar-cleaving enzyme, neuraminidase, coronaviruses must find a balance through modifying glycoprotein binding^49^. Losing domain 0 could therefore be a trade-off to enhance virus release. This may be compensated by the earlier discussed enhanced flexibility of S and increased binding efficiency at the RBD through mutation. These are manifold in the FCoV-23 pCCoV recombinant and in particular mutations at position 546 (already possibly associated with FIP) and 595 are likely to have a strong impact on aminopeptidase N receptor binding (Figure 4C).

Recombination between FCoV-1 and pCCoVs is not surprising in that i) such recombinations have been observed before, and ii) pCCoVs have evolved in Greece and south eastern Europe^35,42^. However, the emergence of pCCoV in Greece happened over 1 years ago so the question as to why this happened now is unclear. One possible explanation is the “right mutation, right time, right place” theory. Recombination between feline and canine coronaviruses happen frequently but as previous reports show, they rarely spread. Introduction of a more successful, spreading variant to a dense population, like the cats in Cyprus, may be sufficient to allow this virus to cause an outbreak. However, the acquisition of a likely in-host deletion adds a further level of concern to coronavirus evolution. These viruses already show a mutation rate, similar to influenza viruses (despite proofreading)^50^, high prevalence for recombination^51^, and here we could show that they are able to acquire significant in-host INDELs that may completely change their cell binding properties and tropism.

This paper reports the emergence of a new *alphacoronavirus 1* feline/canine coronavirus that shows high spreading potential with the associated pathology of lethal FIP if left untreated. Further investigations into the properties of this new virus are now essential. Whilst antivirals, including GS-441524 and molnupiravir were successfully used to treat many cats affected by FCoV-23, early treatment is essential but associated costs can be prohibitive. Prevention of spread and the development of new vaccines are important to stop this epizootic virus from spreading beyond Cyprus.

## Acknowledgements

The authors like to thank the Pancyprian Veterinary Association and all the veterinarians in Cyprus for sample submission. We would like to thank Dr Sam Lycett, The Roslin Institute, The University of Edinburgh for advice on sequence analysis.

## Author contributions

Conceptualization, C.A., A.W., D.G-M., S. M. and C.T-B.; Methodology, C.A., A.W., S.M., and C.T-B.; Validation, C.A., A.W., S.M. and C.T-B.; Formal Analysis, A.W., S.M. and C.T-B.; Investigation, C.A., A.W., D.E., A.H., M.O., S.F., A.M., M.L., R.H., C.T-B.; Resources, C.A., A.W., D.E., M.L., A.H., A.Z., S.L., M.G. and C.T-B.; Data Curation, A.W., S.M., and C. T-B.; Writing – Original Draft Preparation, C.A., A.W., and C.T.B.; Writing – Review & Editing, C.A., A.W., S.F., D.G-M., S.M. and C.T-B.; Visualization, A.W., S.M. and C.T-B.; Supervision, C.A. and C.T-B.; Project Administration, C.A. and C.T-B.; Funding Acquisition, C.A., and C.T-B.

## Methods

### Enrolment of FIP cases in Cyprus

The electronic records of Vet Dia Gnosis Ltd in Limassol, Cyprus (at present, the only Veterinary diagnostics lab on the island), were searched for any positive FCoV RT-PCR or IHC cases from September 2021 up to June 2024. The cases to be enrolled needed to have compatible clinicopathological findings for FIP as recently outlined by the European Advisory Board on Cat Diseases Guidelines^25^ as well having a positive FCoV IHC (on tissue biopsies with granulomatous lesions) or a positive RT-PCR for FCoV in one of the following samples: peritoneal fluid, pleural fluid, cerebrospinal fluid and granuloma fine needle aspiration biopsies or tissue biopsies. The original samples were submitted to the Vet Dia Gnosis Ltd commercial laboratory (Limassol, Cyprus) by local veterinarians and then submitted to LABOKLIN commercial laboratory (Bad Kissingen, Germany). Ethical approval for this study was granted by the Pancyprian Vet Association. According to the terms and conditions of the Vet Dia Gnosis, as well as the Cypriot legislation [The Dogs LAW, N. 184 (I)/2002], no special permission from animal owners or the animal welfare commission is needed for additional testing on residual sample material once diagnostics are completed. According to the terms and conditions of the LABOKLIN laboratory, as well as the RUF-55.2.2.2532-1-86-5 decision of the government of Lower Franconia, no special permission from animal owners or the animal welfare commission is needed for additional testing on residual sample material once diagnostics are completed. The study was also approved by the Veterinary Ethical Review Committee, The Royal (Dick) School of Veterinary Studies (R(D)SVS), The University of Edinburgh, UK (VERC Reference: 233.23).

### Sample from cat imported to the UK

Two UK veterinary practices contacted the R(D)SVS with suspected cases of FIP in cats recently imported from Cyprus. Peritoneal fluid samples were taken from the cats and sent to R(D)SVS for further testing and for sequencing at The Roslin Institute with the consent of the cats’ owners. The veterinary practices and the cats’ owners were kept informed at all stages.

### Histopathology and immunohistochemistry

A sub-set of cases (n=4) from 17 non-effusive (dry) FIP cats diagnosed after January 2023 with available tissue specimens were selected. Tissue specimens were carefully obtained and immediately fixed in 10% buffered formalin. Following fixation, the tissues were processed by embedding, 4μm sectioning, and subsequent staining with hematoxylin and eosin (H&E). The inclusion criteria were defined based on comprehensive histopathological assessments. Specifically, emphasis was placed on identifying the characteristic FIP-associated histological hallmarks, which encompassed vasculitis, phlebitis, and periphlebitis. Consecutive tissue sections were mounted onto charged slides. Following pre-treatment for 5 min at 110°C in 0.01M pH 6 citrate buffer, slides were incubated for 30 min at room temperature with primary mouse monoclonal anti-feline coronavirus antibody at 1:400 (BioRad, MCA 2194). The EnVision anti-mouse system was used for visualization according to the manufacturer’s instructions (Agilent).

### RNA extraction and cDNA synthesis

RNA from specimens from Cyprus underwent automated total nucleic acid extraction using the MagNA Pure 96 DNA AND Viral NA Small Volume kit (Roche Diagnostics). RT-PCR for FCoV was performed at Laboklin Bad Kissingen, Germany^52^.

RNA samples from cats imported to the UK from Cyprus were extracted using the QIAamp viral RNA extraction kit (Qiagen) according to the manufacturer’s instructions.

cDNA synthesis on all RNA samples was carried out using LunaScript RT SuperMix Kit (NEB) with 16 µl template RNA in 20 µl reactions according to the manufacturer’s instructions.

### Primer design and PCR amplification

A set of primers for 1 kb /100 bp overlap tiled amplification were initially designed with available FCoV genomes on NCBI^53^ using primalscheme^54^, and by manual redesign (Supplementary Table S19). These were used to generate a composite representative FCoV-23 genome. Using this genome a new, 800bp/80 bp overlap primer scheme was generated to sequence FCoV-23. The primer sequences used here are found in (Supplementary Table S20). The PCRs for these multiplexed amplifications used the same conditions described below, but with multiplexed primers.

For a subset of samples, DNA was amplified using primers only targeting parts of spike, ORF1b, and ORF3c/E/M. These primers may be found in (Supplementary Table S19, first four rows).

cDNA synthesis was performed using LunaScript reverse transcriptase Supermix (NEB) followed by PCR amplification with Q5 (NEB). 1.25μl 10μM forward and reverse primers and 3μl cDNA in a 25μl reaction. Amplification was performed with the following PCR conditions: initial denaturation at 98°C for 30 seconds, 40 cycles of denaturation at 98°C for 10 seconds, annealing and extension at 65°C for 4 minutes, followed by a final extension at 65°C for 5 minutes. Amplified DNA was purified using AMPure XP beads (Beckman Coulter).

### Nanopore sequencing

Amplicons were quantified using Qubit (Thermo Fisher) High Sensitivity assays and diluted to 150ng DNA per sample in 12μl nuclease-free water. Libraries were prepared using Oxford Nanopore Technologies’ (ONT) NBD112.96 ligation kit following the manufacturer’s protocol for amplicon sequencing with some modifications. Due to unavailability of NBD112.96 kit reagent AMII H, following ligation of barcodes a second end-prep was carried out using the Ultra II End Repair module (NEB). The rest of the protocol was carried out from the adapter ligation stage as per manufacturer’s protocol for LSK112 using the AMX-F adapter supplied in the early access Q20+ version of kit 112. The library was loaded onto a MinION R10.4 flow cell and sequencing was carried out on a GridION sequencing device.

In order to identify the virus present in one of the UK cases as quickly as possible, only the spike amplicon was amplified and the sample sequenced using ONT RAD004 rapid sequencing kit on an R9.4 flow cell following manufacturer’s protocol. The other UK case was amplified at an earlier time and was included in a sequencing run with the Cypriot samples, the data from that sample was treated in the same way as the Cypriot samples and was the sample later found to have the non-recombinant FCoVI spike.

### Bioinformatic analysis

Basecalling and demultiplexing was carried out on the GridION sequencing device (ONT) using Guppy (v2.24-r1122) on super accurate mode, specifying --require_barcodes_both_ends and using appropriate super accurate basecalling models for each of the different sequencing methods used. For the spike amplicons, following basecalling, amplicon_sorter^55^ (v2023-06-19) was used to identify consensus amplicons between 3kb and 5kb without using a reference. Spike amplicons were identified from the output via alignment with minimap2^56^ (v2.22) to the spike from the first sample we sequenced (1-G7_Gi_6739) for which a consensus was made using the same process, with the correct amplicon identified via BLAST^57^. The identified amplicons were polished with medaka (v1.8.0, ONT). For the UK case which was sequenced with the rapid kit, because the reads were fragmented by the library prep process, reads were used to polish the 1-G7_Gi_6739 sequence using medaka. The reads were aligned to the polished sequence with minimap2 and visualized with IGV^58^ (v2.11.1) to visually confirm the reads supported the consensus sequence and that it had not been biased by the reference used. The amplicons in POL1ab and Orf3c/E/M were assembled and polished using LILO^59^. Multisequence alignments were carried out for each of the regions using mafft^60^ (v7.49) against all complete genomes of CCoV and FCoV genomes available on NCBI using the --adjustdirection flag. Alignments were visualized in Mega7^61^. All downloaded genomes were trimmed down to each of the target regions amplified from our samples. Maximum likelihood trees were constructed using IQ-TREE^62^ (v2.0.5). TempEst^63^ (v1.5.3) was used to determine the best fitting root for the trees, and visualizations and annotations of the trees were done using iTOL^64^ (v6.8.1).

To assemble a representative genome for the population, the combined reads of several samples for which we had sequencing data were run through LILO. The polished amplicons were mapped to a single fasta file containing an FCoVI (MT239440.1) and a CCoVII (KP981644.1) using minimap2 and visualized with IGV, with only the amplicons from the spike region aligning to CCoVII. Representative amplicons covering the entire genome were selected and scaffolded using scaffold_builder^65^ with MT239440.1 as a reference. The raw reads were trimmed by 50bp to remove adapters and barcodes and they were used to polish this scaffold using medaka. Mafft was used to align this genome to representative genomes from *Alphacoronavirus 1* with Canine Respiratory Coronavirus as an outgroup, and Mega7 was used to create a neighbor join tree which was visualized and annotated with iTOL.

Having assembled a representative genome, primers were redesigned to include ambiguous bases that supported the sequence in the representative genome, and these are the primers that were used to amplify 800bp tiled amplicons for individual Cypriot samples. The Cypriot samples were assembled using LILO, with the representative genome as the reference. Assembled genomes had their reads aligned back to them using minimap2 and were visualised with IGV. Each genome was visually inspected for errors and corrections made, primarily where primer sequences had been incorrectly incorporated. Where there were gaps in these genomes, and there were reads that spanned the gap but did not reach the depth threshold for LILO, reads were extracted and amplicon_sorter was used to attempt to create a consensus for the missing amplicon. These amplicons were carefully visually inspected for errors before being incorporated into the assembly to close gaps using scaffold_builder. Multisequence alignments between the population level representative genome. a sample without a spike domain 0 deletion (2-C11_Re_10276) and a sample with a domaion 0 deletion (2-F12_Bl_11350) against MT239440.1, LC742526.1 and KP981644.1 was carried out using ClustalW^66^ in Mega7, and recombination analyses carried out using RDP5^37^ (v.5.45) on default settings.

### Structure prediction of spike deletion variants

SWISS-MODEL structure prediction and analysis (https://swissmodel.expasy.org/, accessed October 2023)^67^ were used to model the partial, high confidence N-terminal sequence of a full-length spike (1008aa) and a deletion spike (797aa). To model the G8 Cypriot full-length spike, we used the alphacoronavirus 1 experimentally resolved CCoV-HuPn-2018^40^ structure as a template. Modelling against the 7usa.1.a, swung out confirmation, yielded GMQE 0.60 and Global QMEAND of 0.68±0.05 with a sequence identity of 88.97% and against 7us6.1.A, proximal confirmation, yielded GMQE of 0.62 and Global QMEAND of 0.58±0.05 with a sequence identity of 81.03%. Modelling of C10 spike, C-terminal deletion, against 7usa.1.A, swung out confirmation, yielded GMQE 0.74 and Global QMEAND of 0.69±0.05 with a sequence identity of 88.66% and against 7us6.1.A, proximal confirmation of HuPN, yielded GMQE of 0.76 and Global QMEAND of 0.70±0.05 with a sequence identity of 88.42%.

The receptor binding domain was modelled using the 7u0I.1B structure, again CCoV-HuPn-2018^40^ complexed with canine APN as a template. GMQE was 0.89 and Global QMEAND 0.85±0.07 with a sequence identity of 92.41%. PDB files are available in the supplementary materials.

