## Supplementary Figures for "Emergence and spread of feline infectious peritonitis due to a highly pathogenic canine/feline recombinant coronavirus"

### Supplementary figure S1

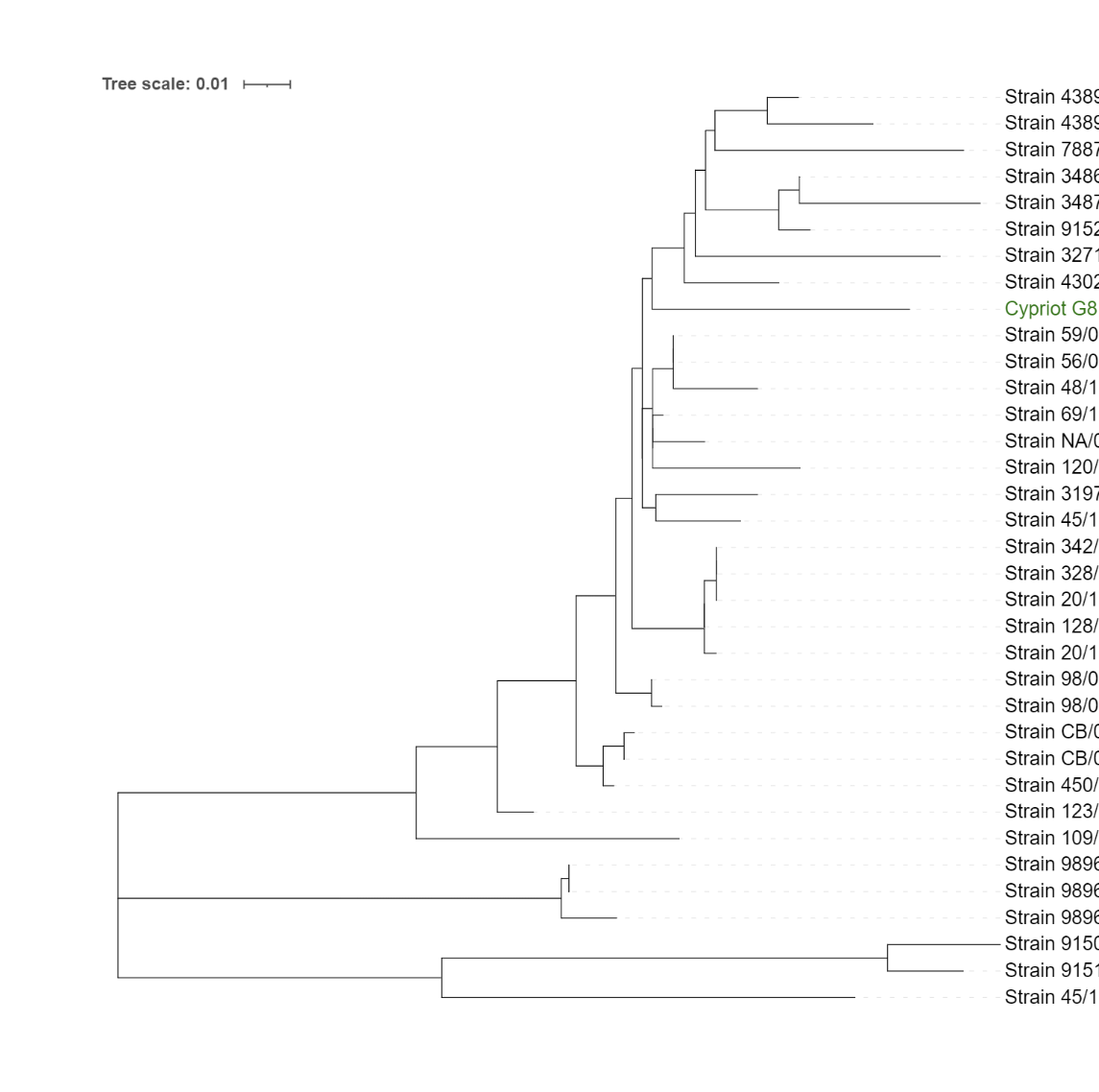

**Supplementary Figure S1: Relationship to other pCCoV viruses.** Maximum likelihood tree of pantropic CCoV spikes (~450bp region). Alignments were done with Muscle in MEGA7, and maximum likelihood tree was made in MEGA7 with default settings. Tree was visualised in iTOL.

Supplementary figure 1 shows a maximum likelihood tree generated from the alignment of known pCCoV spike amplicons with the non-deletion form of the FCoV-23 amplicon. The alignment was carried out with Muscle^1^ in MEGA7^2^ and the maximum likelihood tree was generated with MEGA7 on default settings. The tree was visualised with iTIOL^3^. The region targeted is ~450bp with longer sequences trimmed down. The FCoV-23 spike with the deletion could not be used as the deleted region heavily overlaps with the region in the alignment. The FCoV-23 amplicon clusters among the pCCoV sequences.

### Supplementary figure S2

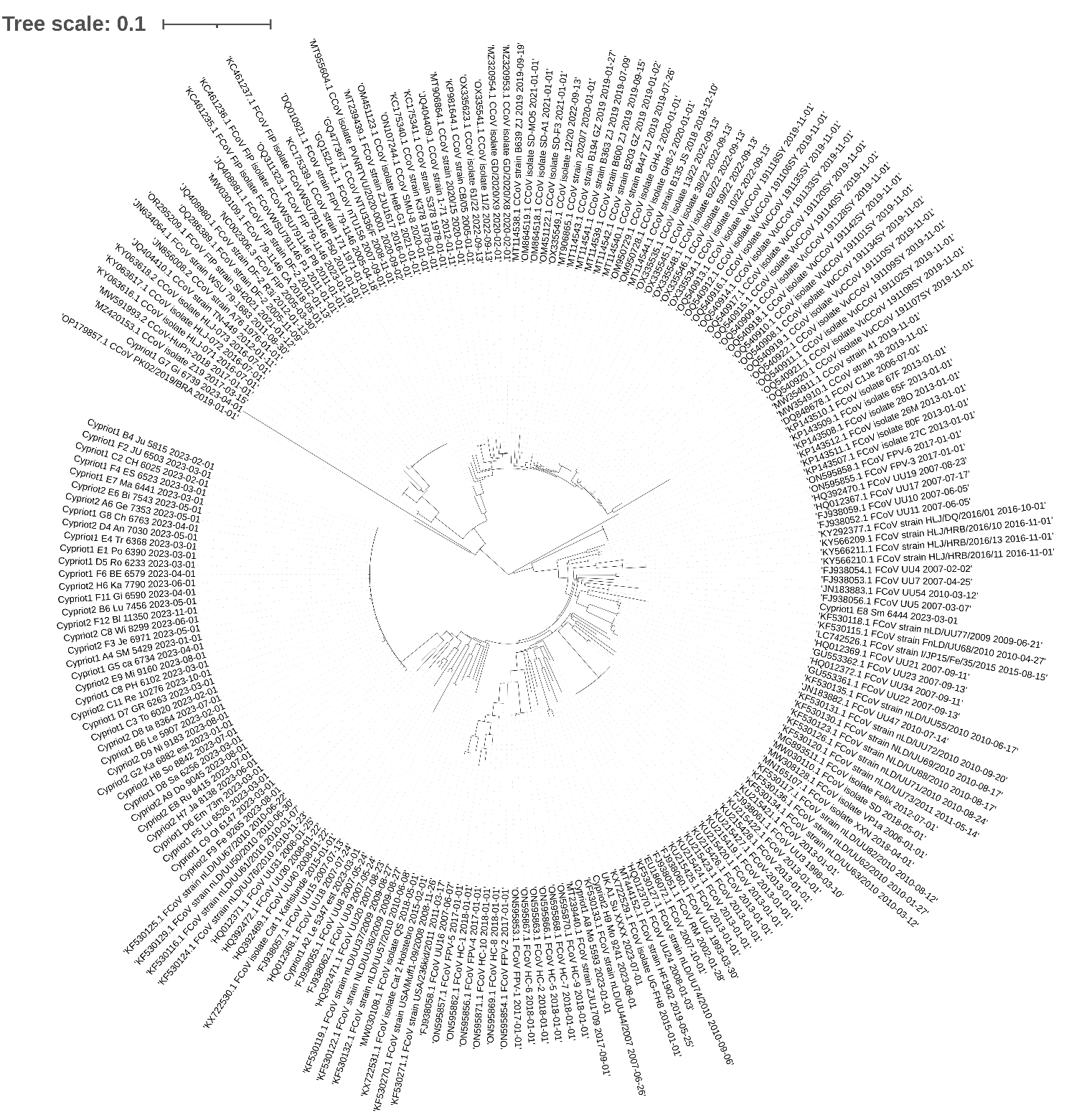

**Supplementary Figure S2: ORF1b ML tree from figure 2.** Higher resolution version of the maximum likelihood tree for a ORF1b region. The amplicon for this region was amplified with the primers listed at the top of supplementary table S19.

### Supplementary figure S3

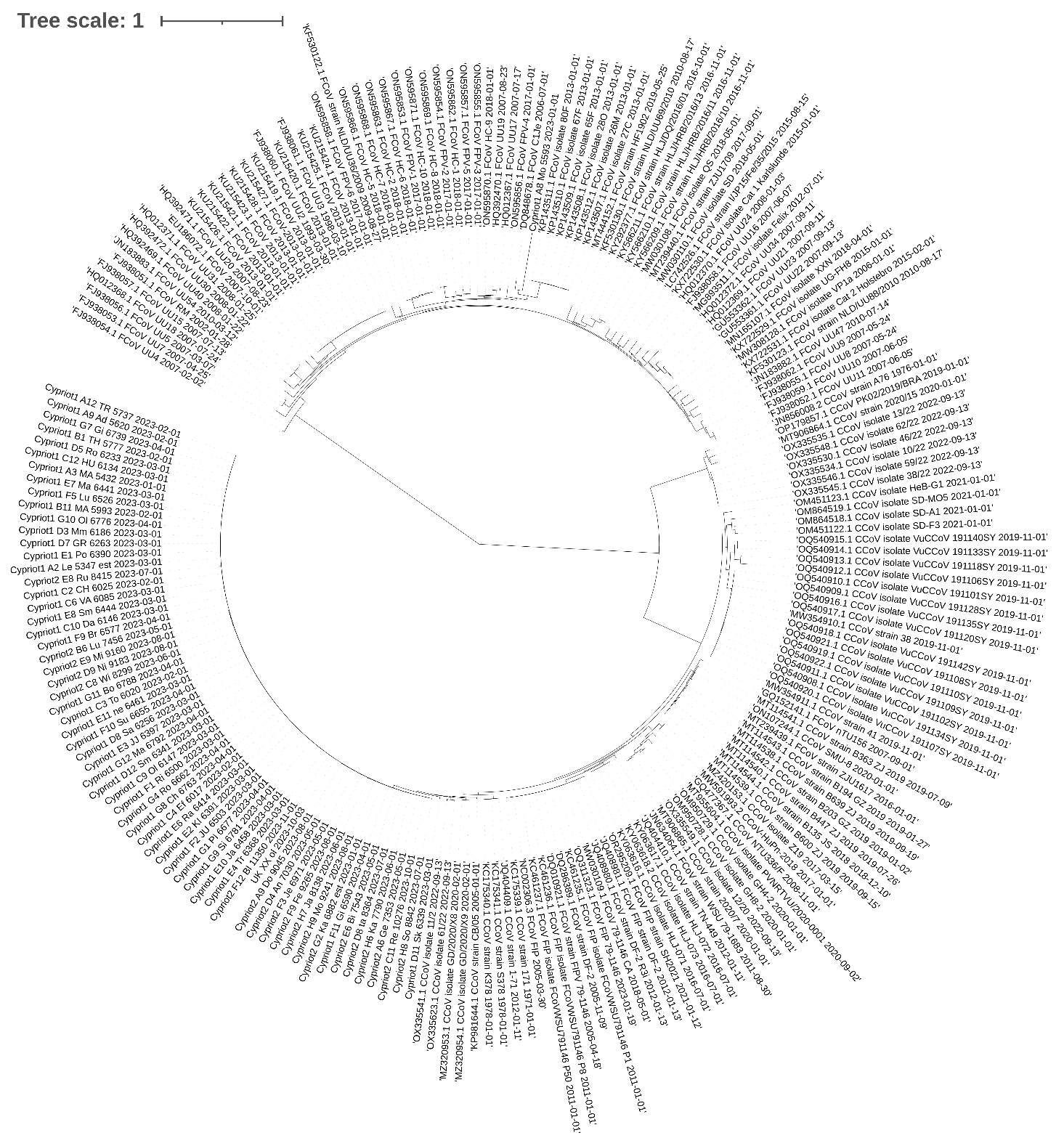

**Supplementary Figure S3: Spike ML tree from figure** **S2.** Larger version of the maximum likelihood tree for the spike gene. The amplicon for this region was amplified with the primers listed at the top of supplementary table S19.

### Supplementary figure S4

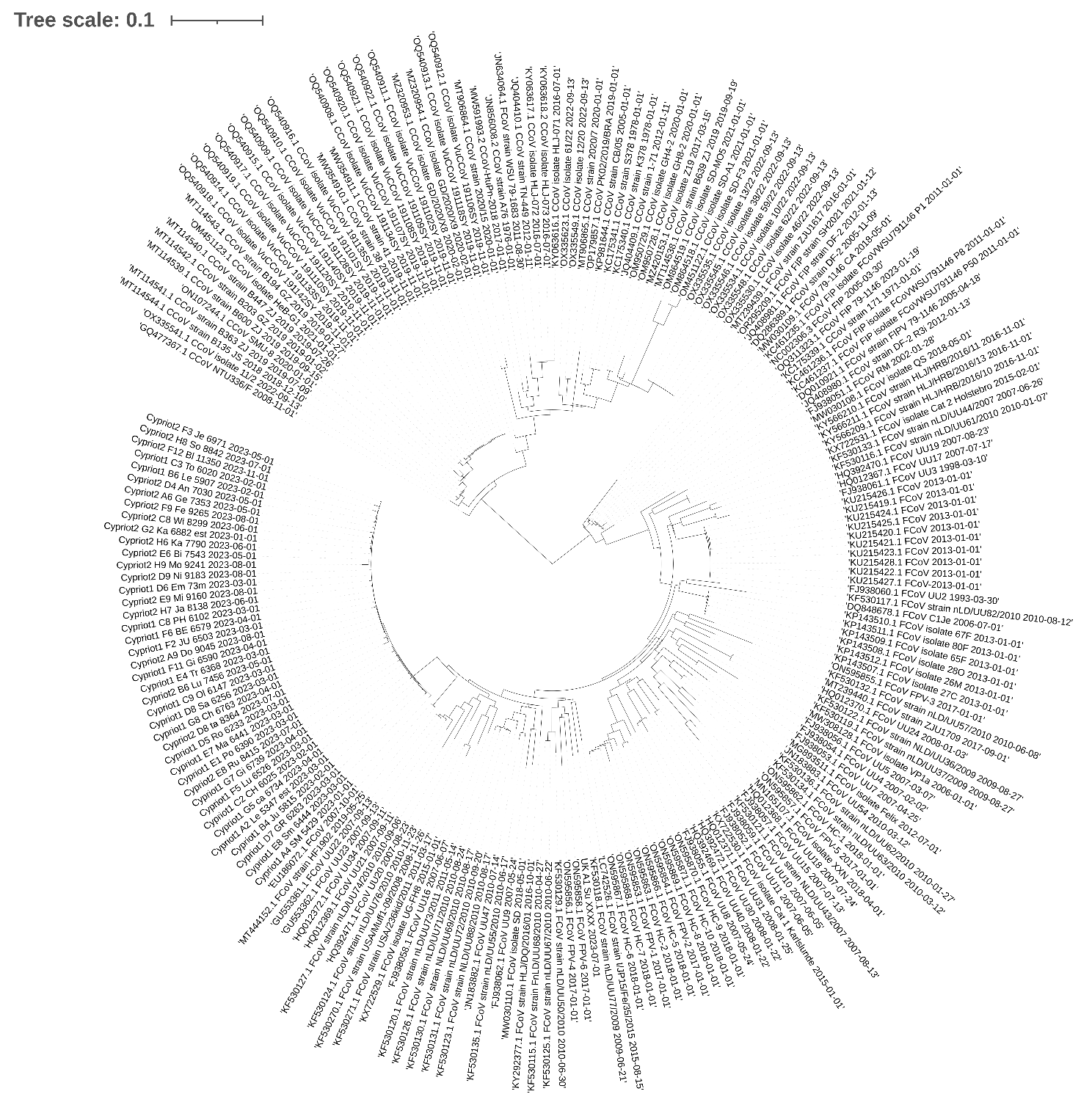

**Supplementary Figure S4: ORF3c/E/M ML tree from figure 2.** Larger version of the maximum likelihood tree for a region spanning the NSP3c, E and M genes. The amplicons for this region were amplified with the primers listed at the top of supplementary table S19.

### Supplementary figure S5

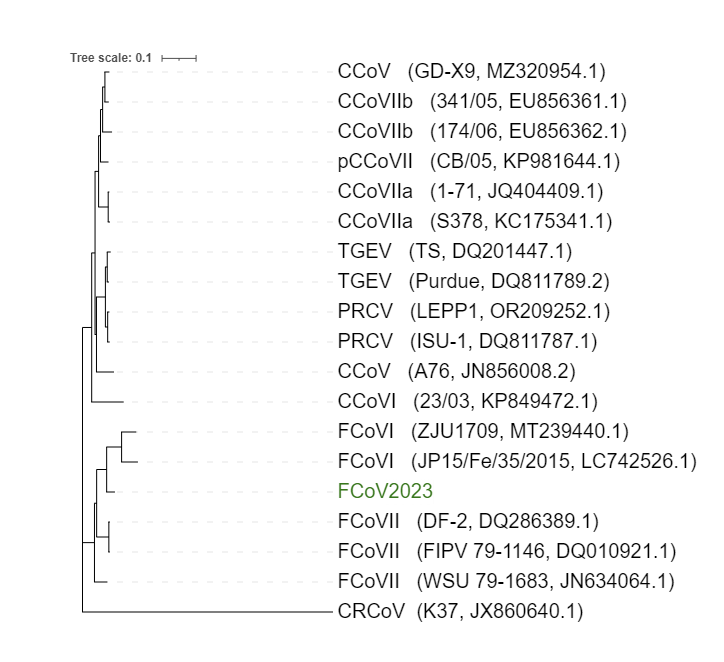

**Supplementary figure S5: Maximum likelihood tree of FCoV-23 with other alphacoronavirus 1 full genome sequences.**

### Supplementary figure S6

**A**
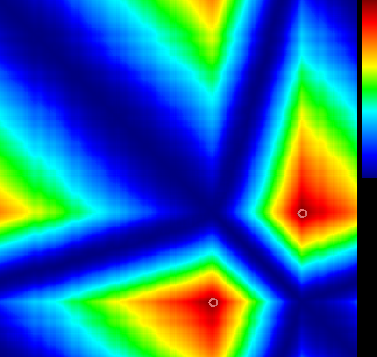

**B**
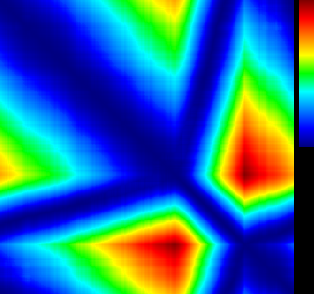

**Supplementary Figure S6: MaxChi**^4^ **breakpoint matrix from RDP5**^5^ **analysis for A) spike full-lenth FCoV-23 genome 2-C11 Re 10276 and B) domain 0-deletion FCoV-23 genome 2-F12 BW 11350.** MaxChi breakpoint matrix generated with default settings in RDP5. Supplementary figure 3 shows the MaxChi7 breakpoint matrix generated in RDP58. The dark red region highlights the likely recombination breakpoints, which align with the breakpoints described in the main text.

### Supplementary figure S7

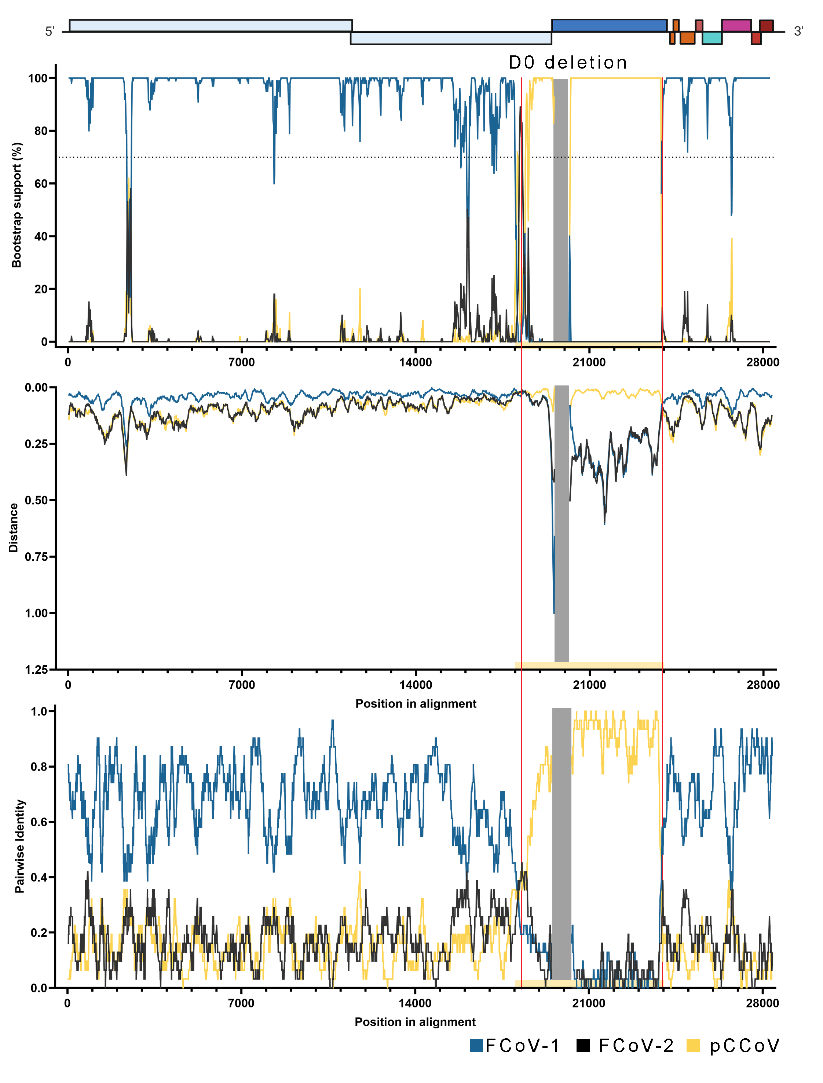

**Supplementary figure S7: Recombination analysis of domain 0-deletion FCoV-23 genome 2-F12 BW 11350.** A) Visualizations of a recombination analysis carried out on the assembled FCoV-23 genome and representative genomes of FCoV-1 (blue), FCoV-2 (black) and pCCoV (yellow). An annotated sequence can be found in Supplementary File 1[genbank]. The yellow panel shows the likely recombination break region, with a red vertical line showing the likely break point. The first panel shows the results of the Bootscan analysis, the second panel shows the sequence distance, and the third panel shows the RDP pairwise identity analysis. All three panels show good support for the recombination between FCoV-1 and pCCoV. These results are further supported by high statistical likelihood of recombination shown in Supplementary Table S22.

### Supplementary figure S8

**A**
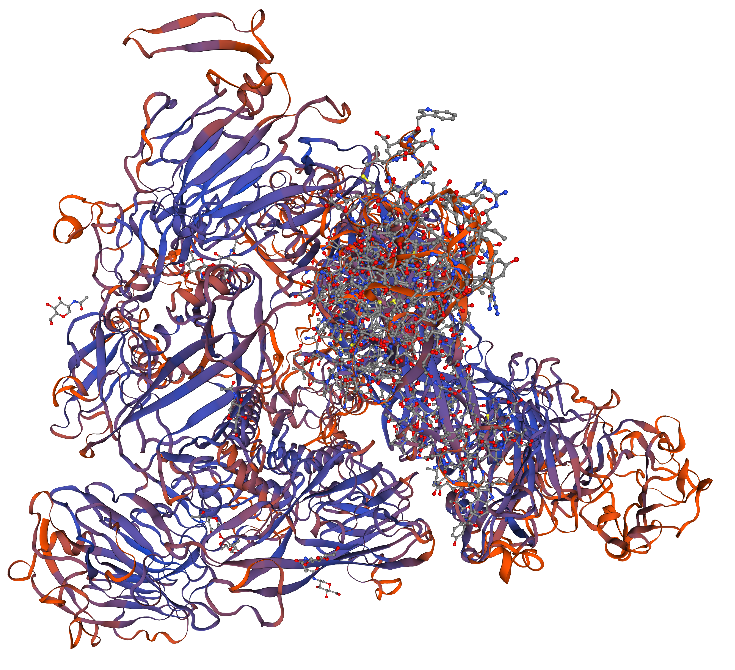
 **B**
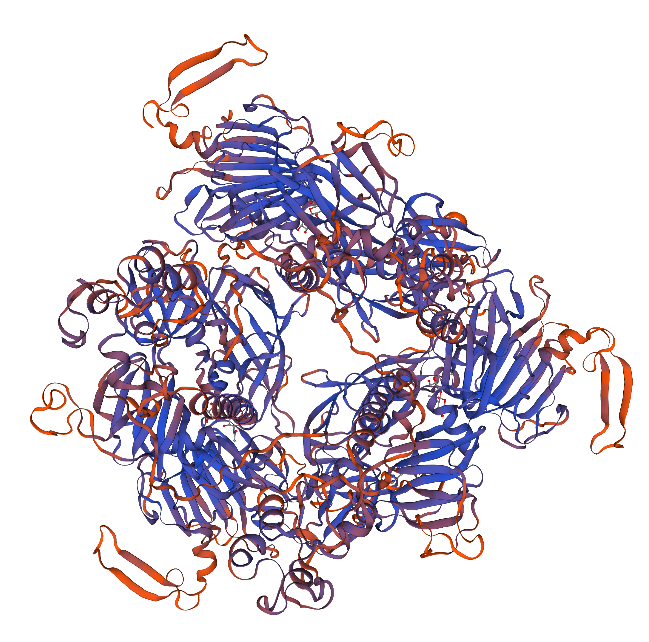

**C**
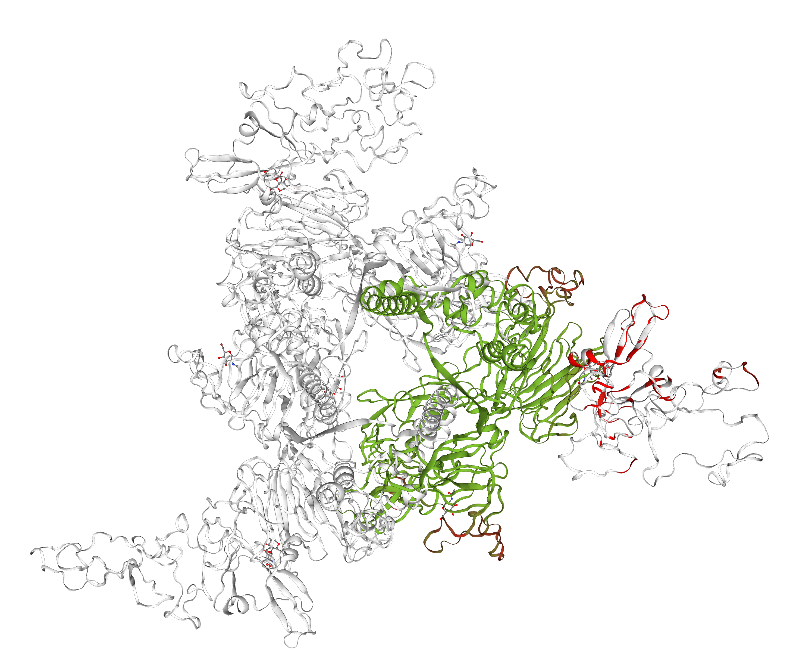
**D**
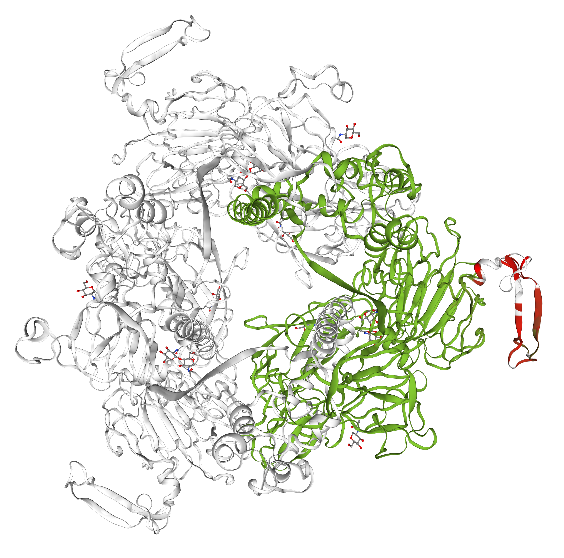

**Supplementary figure S8: Protein structure modelling.** Structural modelling of A) G8 full-length and B) C10 domain 0 deletion (short) spike using the 7us6.1.A proximal confirmation ofCCoV-HuPN-2018 as a template. C&D) Comparison between the G8 full-length and the C10 domain 0 deletion (short) spike structure prediction. C) represents the swung out (modelled against the 7usa.1.A template) and D) the proximal confirmation (modelled against the 7us6.1.A template). Colors represent consistency with red being inconsistent and green consistent between the two structures.

### Supplementary figure S9

|  | **1-F11** | **2-A6** | **2-A9** | **2-B7** | **2-C11** | **2-C8** | **2-D4** | **2-D8** | **2-D9** | **2-E6** | **2-E8** | **2-E9** | **2-F12** | **2-F3** | **2-F9** | **2-G2** | **2-H6** | **2-H7** | **2-H8** | **2-H9** |
| --- | --- | --- | --- | --- | --- | --- | --- | --- | --- | --- | --- | --- | --- | --- | --- | --- | --- | --- | --- | --- |
| **1-F11** |  | 0.99412 | 0.99438 | 0.99739 | 0.99718 | 0.99675 | 0.99722 | 0.99831 | 0.99739 | 0.99351 | 0.96161 | 0.99661 | 0.99604 | 0.99467 | 0.99682 | 0.99817 | 0.99566 | 0.99400 | 0.99619 | 0.95883 |
| **2-A6** | 0.00588 |  | 0.99388 | 0.99314 | 0.99288 | 0.99398 | 0.99292 | 0.99369 | 0.99292 | 0.99545 | 0.96153 | 0.99446 | 0.99193 | 0.99470 | 0.99277 | 0.99383 | 0.99600 | 0.99592 | 0.99211 | 0.95894 |
| **2-A9** | 0.00562 | 0.00612 |  | 0.99372 | 0.99312 | 0.99417 | 0.99393 | 0.99361 | 0.99311 | 0.99312 | 0.95927 | 0.99432 | 0.99217 | 0.99412 | 0.99339 | 0.99383 | 0.99503 | 0.99358 | 0.99218 | 0.95743 |
| **2-B7** | 0.00261 | 0.00686 | 0.00628 |  | 0.99617 | 0.99631 | 0.99674 | 0.99678 | 0.99624 | 0.99263 | 0.96203 | 0.99563 | 0.99541 | 0.99377 | 0.99624 | 0.99690 | 0.99447 | 0.99302 | 0.99509 | 0.95893 |
| **2-C11** | 0.00282 | 0.00712 | 0.00688 | 0.00383 |  | 0.99548 | 0.99597 | 0.99644 | 0.99573 | 0.99283 | 0.96027 | 0.99637 | 0.99484 | 0.99330 | 0.99563 | 0.99661 | 0.99478 | 0.99260 | 0.99506 | 0.95757 |
| **2-C8** | 0.00325 | 0.00602 | 0.00583 | 0.00369 | 0.00452 |  | 0.99604 | 0.99605 | 0.99569 | 0.99308 | 0.96052 | 0.99576 | 0.99448 | 0.99396 | 0.99591 | 0.99634 | 0.99509 | 0.99357 | 0.99463 | 0.95764 |
| **2-D4** | 0.00278 | 0.00708 | 0.00607 | 0.00326 | 0.00403 | 0.00396 |  | 0.99647 | 0.99601 | 0.99226 | 0.96091 | 0.99543 | 0.99507 | 0.99341 | 0.99618 | 0.99672 | 0.99442 | 0.99276 | 0.99497 | 0.95995 |
| **2-D8** | 0.00169 | 0.00631 | 0.00639 | 0.00322 | 0.00356 | 0.00395 | 0.00353 |  | 0.99669 | 0.99297 | 0.96124 | 0.99598 | 0.99548 | 0.99408 | 0.99619 | 0.99747 | 0.99513 | 0.99361 | 0.99559 | 0.95818 |
| **2-D9** | 0.00261 | 0.00708 | 0.00689 | 0.00376 | 0.00427 | 0.00431 | 0.00399 | 0.00331 |  | 0.99217 | 0.96047 | 0.99548 | 0.99491 | 0.99352 | 0.99559 | 0.99724 | 0.99441 | 0.99270 | 0.99502 | 0.95745 |
| **2-E6** | 0.00649 | 0.00455 | 0.00688 | 0.00737 | 0.00717 | 0.00692 | 0.00774 | 0.00703 | 0.00783 |  | 0.95991 | 0.99454 | 0.99117 | 0.99645 | 0.99188 | 0.99312 | 0.99534 | 0.99632 | 0.99137 | 0.95613 |
| **2-E8** | 0.03839 | 0.03847 | 0.04073 | 0.03797 | 0.03973 | 0.03948 | 0.03909 | 0.03876 | 0.03953 | 0.04009 |  | 0.96041 | 0.95982 | 0.96054 | 0.96010 | 0.96535 | 0.95980 | 0.96002 | 0.95958 | 0.95349 |
| **2-E9** | 0.00339 | 0.00554 | 0.00568 | 0.00437 | 0.00363 | 0.00424 | 0.00457 | 0.00402 | 0.00452 | 0.00546 | 0.03959 |  | 0.99434 | 0.99504 | 0.99499 | 0.99623 | 0.99624 | 0.99438 | 0.99442 | 0.95761 |
| **2-F12** | 0.00396 | 0.00807 | 0.00783 | 0.00459 | 0.00516 | 0.00552 | 0.00493 | 0.00452 | 0.00509 | 0.00883 | 0.04018 | 0.00566 |  | 0.99277 | 0.99459 | 0.99571 | 0.99339 | 0.99193 | 0.99406 | 0.95708 |
| **2-F3** | 0.00533 | 0.00530 | 0.00588 | 0.00623 | 0.00670 | 0.00604 | 0.00659 | 0.00592 | 0.00648 | 0.00355 | 0.03946 | 0.00496 | 0.00723 |  | 0.99292 | 0.99444 | 0.99542 | 0.99560 | 0.99266 | 0.95733 |
| **2-F9** | 0.00318 | 0.00723 | 0.00661 | 0.00376 | 0.00437 | 0.00409 | 0.00382 | 0.00381 | 0.00441 | 0.00812 | 0.03990 | 0.00501 | 0.00541 | 0.00708 |  | 0.99646 | 0.99427 | 0.99242 | 0.99471 | 0.95753 |
| **2-G2** | 0.00183 | 0.00617 | 0.00617 | 0.00310 | 0.00339 | 0.00366 | 0.00328 | 0.00253 | 0.00276 | 0.00688 | 0.03465 | 0.00377 | 0.00429 | 0.00556 | 0.00354 |  | 0.99545 | 0.99383 | 0.99586 | 0.96322 |
| **2-H6** | 0.00434 | 0.00400 | 0.00497 | 0.00553 | 0.00522 | 0.00491 | 0.00558 | 0.00487 | 0.00559 | 0.00466 | 0.04020 | 0.00376 | 0.00661 | 0.00458 | 0.00573 | 0.00455 |  | 0.99534 | 0.99394 | 0.95669 |
| **2-H7** | 0.00600 | 0.00408 | 0.00642 | 0.00698 | 0.00740 | 0.00643 | 0.00724 | 0.00639 | 0.00730 | 0.00368 | 0.03998 | 0.00562 | 0.00807 | 0.00440 | 0.00758 | 0.00617 | 0.00466 |  | 0.99169 | 0.95640 |
| **2-H8** | 0.00381 | 0.00789 | 0.00782 | 0.00491 | 0.00494 | 0.00537 | 0.00503 | 0.00441 | 0.00498 | 0.00863 | 0.04042 | 0.00558 | 0.00594 | 0.00734 | 0.00529 | 0.00414 | 0.00606 | 0.00831 |  | 0.95650 |
| **2-H9** | 0.04117 | 0.04106 | 0.04257 | 0.04107 | 0.04243 | 0.04236 | 0.04005 | 0.04182 | 0.04255 | 0.04387 | 0.04651 | 0.04239 | 0.04292 | 0.04267 | 0.04247 | 0.03678 | 0.04331 | 0.04360 | 0.04350 |  |
|  |  | Mean | Min | Max |  |  |  |  |  |  |  |  |  |  |  |  |  |  |  |  |
|  | All | 0.4487 | 0.9975 | 0.0017 |  |  |  |  |  |  |  |  |  |  |  |  |  |  |  |  |
|  | "Exclude |  |  |  |  |  |  |  |  |  |  |  |  |  |  |  |  |  |  |  |
|  | H9&E8" | 0.2224 | 0.0017 | 0.9968 |  |  |  |  |  |  |  |  |  |  |  |  |  |  |  |  |

**Supplementary Figure S9: Sequence identity / divergence of FCoV-23 full genome sequences.** The number of base substitutions / identical bases per site from between sequences are shown. Analyses were conducted using the Maximum Composite Likelihood model^6^ using 20 full-genome FCoV-23 sequences using 29,144 base positions. All Codon positions (1/2/3 +NC) were included. Ambiguous positions were removed using the pairwise deletion option. Evolutionary analyses were conducted in MEGA X^7^ and are displayed (bottom) with or without sequences H9 and E8, which display poorer sequence quality (lower coverage and bigger gaps) and should be treated with caution.

### Supplementary figure S10

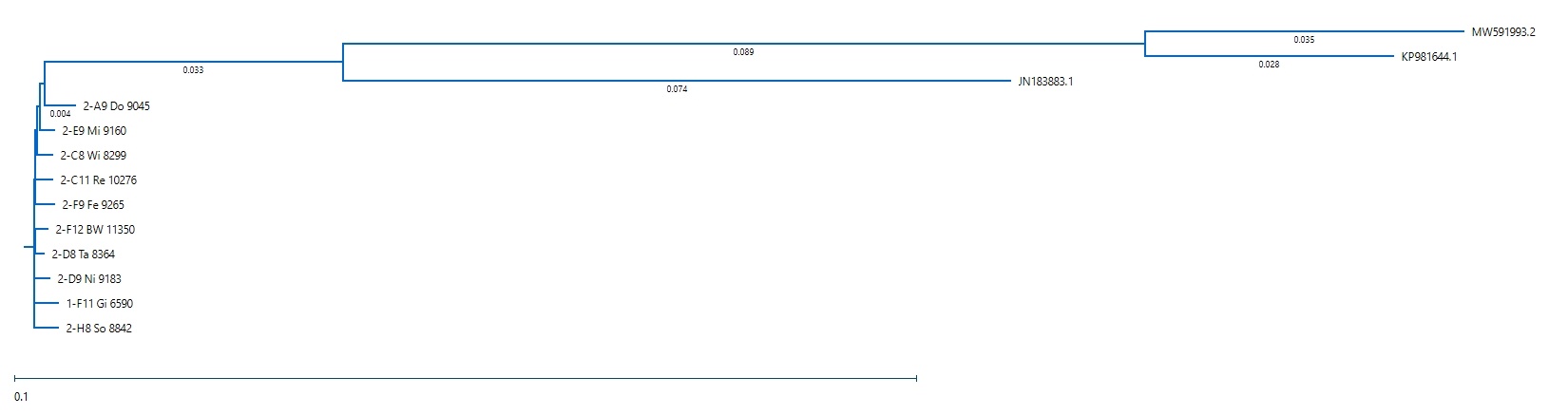

**Supplementary figure S10: Clustal W whole genome alignment.** 10 gap-free FCoV-23 genome sequences were multi-genome aligned using ClustalW with JN183883 (closest FCoV-2, UU54), KP981644 (pantropic CCoV CB/05), and MW591993 (CCoV-HuPn-2018) for context.
