## Supplementary Tables for "Emergence and spread of feline infectious peritonitis due to a highly pathogenic canine/feline recombinant coronavirus"

**Supplementary table S1: FIP outbreak Cyprus Jan 2023-Jun 2024; case distribution – sex**

| Sex | n | Total | Percentage |
| --- | --- | --- | --- |
| Female | 65 | 215 | 33.02 |
| Male | 137 | 215 | 66.98 |

**Supplementary table S2: FIP outbreak Cyprus Jan 2023-Jun 2024;
case distribution – Age stats (years); the age of 51 cases are missing**

|  | Mean_age | Median_age | Min_age | Max_age |
| --- | --- | --- | --- | --- |
| Jan 23- Mar-24 | **4.6059** | 3 | 1 | 19 |
| Jan 23- Sep-23 | **4.4127** | 3 | 1 | 16 |
| Oct-23-Mar-24 | **5.5423** | 4.25 | 1 | 19 |
| Apr-24-Jun-24 | **5.1250** | 3.7 | 0.6 | 13.3 |

**Supplementary table S3: FIP outbreak Cyprus Jan 2023 -Jun 2024;
case distribution – FIP form**

| Form | n | Total | Percentage |
| --- | --- | --- | --- |
| Non-effusive | 22 | 215 | 10.23 |
| Neurological | 56 | 215 | 26.05 |
| Effusive | 137 | 215 | 63.72 |

**Supplementary table S4: FIP outbreak Cyprus Jan 2023 -Sep 2023;
case distribution – FIP form**

| Form | n | Total | Percentage |
| --- | --- | --- | --- |
| Non-effusive | 4 | 167 | 2.4 |
| Neurological | 48 | 167 | 28.74 |
| Effusive | 115 | 167 | 68.86 |

**Supplementary table S5A: FIP outbreak Cyprus Oct 2023 -Mar 2024;
case distribution – FIP form**

| Form | n | Total | Percentage |
| --- | --- | --- | --- |
| Non-effusive | 12 | 34 | 35.29 |
| Neurological | 8 | 34 | 23.53 |
| Effusive | 14 | 34 | 41.18 |

**Supplementary table S5B: FIP outbreak Cyprus Apr 2024 -Jun 2024;
case distribution – FIP form**

| Form | n | Total | Percentage |
| --- | --- | --- | --- |
| Non-effusive | 6 | 14 | 42.86 |
| Neurological | 0 | 14 | 0 |
| Effusive | 8 | 14 | 57.14 |

**Supplementary table S6: FIP outbreak Cyprus Jan 2023-Mar 2024;
case distribution – District**

| District | n | Total | Percentage |
| --- | --- | --- | --- |
| Nicosia | 58 | 215 | 26.98 |
| Famagusta | 52 | 215 | 24.19 |
| Larnaca | 41 | 215 | 19.07 |
| Limassol | 34 | 215 | 15.81 |
| Paphos | 30 | 215 | 13.95 |

**Supplementary table S7: RNA samples for sequencing; case distribution – Sex**

| Sex | n | Total | Percentage |
| --- | --- | --- | --- |
| Female | 54 | 163 | 33.13 |
| Male | 109 | 163 | 66.87 |

**Supplementary table S8: RNA samples for sequencing; case distribution – Age stats (years).** The ages of 38 cats are missing.

| Mean_age | Median_age | Min_age | Max_age |
| --- | --- | --- | --- |
| 4.21 | 3 | 1 | 16 |

**Supplementary table S9: RNA samples for sequencing; case distribution – FIP form**

| Form | n | Total | Percentage |
| --- | --- | --- | --- |
| Non-effusive | 6 | 163 | 3.68 |
| Neurological | 42 | 163 | 25.77 |
| Effusive | 113 | 163 | 70.55 |

**Supplementary table S10: RNA samples for sequencing; case distribution – District**

| District | n | Total | Percentage |
| --- | --- | --- | --- |
| Nicosia | 46 | 163 | 28.22 |
| Famagusta | 46 | 163 | 28.22 |
| Larnaca | 31 | 163 | 19.02 |
| Limassol | 21 | 163 | 12.88 |
| Paphos | 17 | 163 | 10.43 |
| UK | 2 | 163 | 1.23 |

**Supplementary table S11: RNA samples for sequencing; case distribution – Collection dates**

| District | n | Total | Percentage |
| --- | --- | --- | --- |
| October-21 | 1 | 163 | 0.61 |
| November-21 | 1 | 163 | 0.61 |
| December-21 | 1 | 163 | 0.61 |
| February-22 | 1 | 163 | 0.61 |
| March-22 | 1 | 163 | 0.61 |
| August-22 | 1 | 163 | 0.61 |
| December-22 | 1 | 163 | 0.61 |
| January-23 | 7 | 163 | 4.29 |
| February-23 | 23 | 163 | 14.11 |
| March-23 | 36 | 163 | 22.09 |
| April-23 | 19 | 163 | 11.66 |
| September-23 | 28 | 163 | 17.18 |
| October-23 | 14 | 163 | 8.59 |
| November-23 | 5 | 163 | 3.07 |
| Unknown | 1 | 163 | 0.61 |

**Supplementary table S12: RNA samples for sequencing; case distribution – Sample type used for extraction**

| District | n | Total | Percentage |
| --- | --- | --- | --- |
| Peritoneal | 90 | 163 | 55.21 |
| Pleural | 24 | 163 | 14.72 |
| CSF | 43 | 163 | 26.38 |
| Nasal Swab | 1 | 163 | 0.61 |
| Tissue/LN^a^ intestinal | 5 | 163 | 3.07 |

^a^LN – lymph node

**Supplementary table S13: Successfully sequenced spike protein; case distribution – Sex**

| Sex | n | Total | Percentage |
| --- | --- | --- | --- |
| Female | 25 | 63 | 39.68 |
| Male | 38 | 63 | 60.32 |

**Supplementary table S14: Successfully sequenced spike protein; case distribution – Age stats (years).** The ages of 14 cats are missing

| Mean_age | Median_age | Min_age | Max_age |
| --- | --- | --- | --- |
| 4.44 | 3 | 1 | 16 |

**Supplementary table S15: Successfully sequenced spike protein; case distribution – FIP form**

| Form | n | Total | Percentage |
| --- | --- | --- | --- |
| Non-effusive | 5 | 63 | 7.94 |
| Neurological | 10 | 63 | 15.87 |
| Effusive | 48 | 63 | 76.19 |

**Supplementary table S16: Successfully sequenced spike protein; case distribution – District**

| District | n | Total | Percentage |
| --- | --- | --- | --- |
| Nicosia | 17 | 63 | 26.98 |
| Famagusta | 24 | 63 | 38.10 |
| Larnaca | 7 | 63 | 11.11 |
| Limassol | 9 | 63 | 14.29 |
| Paphos | 4 | 63 | 6.35 |
| UK | 2 | 63 | 3.17 |

**Supplementary table S17: Successfully sequenced spike protein; case distribution – Collection dates**

| Date | n | Total | Percentage |
| --- | --- | --- | --- |
| January-23 | 3 | 63 | 4.76 |
| February-23 | 8 | 63 | 12.70 |
| March-23 | 20 | 63 | 31.75 |
| April-23 | 12 | 63 | 19.05 |
| May-23 | 5 | 63 | 7.94 |
| June-23 | 3 | 63 | 4.76 |
| July-23 | 3 | 63 | 4.76 |
| August-23 | 5 | 63 | 7.94 |
| September-23 | 1 | 63 | 1.59 |
| October-23 | 2 | 63 | 3.17 |
| November-23 | 2 | 63 | 3.17 |

**Supplementary table S18: Successfully sequenced spike protein; case distribution – Sample type used for extraction**

| Form | n | Total | Percentage |
| --- | --- | --- | --- |
| Peritoneal | 41 | 63 | 65.08 |
| Pleural | 8 | 63 | 12.70 |
| CSF | 10 | 63 | 15.87 |
| Tissue/LN^a^ intestinal | 4 | 63 | 6.35 |

**^a^**LN – lymph node

**Supplementary table S19: Preliminary primer sequences used during first amplifications of the FCoV-23 genome.** Primers were designed either using primal scheme^1^ or through manual design following multi-sequence alignment of available FCoV whole genomes on NCBI^2^ with Mafft^3^ (v7.490). This is not a finalised scheme and Supplementary table 20 primers should be used for sequencing FCoV-23.

| Scheme location | Forward (5'-3') | Reverse (5'-3') | Specific target (where applicable) | Start (consensus genome) | End (consensus genome) | Expected length | Note |
| --- | --- | --- | --- | --- | --- | --- | --- |
| 28 to 32 | GACGCAGACTTCAGTGTTA | ACCATTATGCCATTRTARTA | Spike | 19728 | 23341 | 3613 | ^a^ |
| 23 | TGCCCAGCTGARATTGTTAARACAG | CATAGTTGTAAGYTCAAGACCACC | ORF1b region | 16376 | 17426 | 1050 |  |
| 35 | TGTCTBAGTACTGGHTGYTGTGG | AGCATAGGGTCTACAAAATGCAAC | Orf3c/E/M region part 1 | 23592 | 24616 | 1024 |  |
| 36 | GATGGCATTGTKACAAYAACTGTCTT | ARCCGAACATTACATATCTGGAAACTT | Orf3c/E/M region part 2 | 24380 | 25489 | 1109 |  |
| 1 | GGACACCAACTCGAACTAAACGA | GTCCARTCACCDACACCACTACT |  | 0 | 992 | 992 |  |
| 2 | GTAGCACCRCCAGTCAAGARRAAYTC | GTRGCRAARAATGCACTATCAAGRCC |  | 801 | 1775 | 974 |  |
| 3 | GTGAARGCHTTYGATGTYTTCACACA | TACRGGCACACTATGCAGTCTTAA |  | 1605 | 2676 | 1071 |  |
| 3 to 4 | TTGTCAAGCTTGTCAGTGT | ATTGAGCATCGTCTCCAA |  | 1705 | 3726 | 2021 |  |
| 5 to 7 | TGGGCTGYTGCTGTYGAYGAACA | GTGTTTAAGYGCGAGAACTGMCTT |  | 3438 | 6408 | 2970 |  |
| 8 to 9 | CAGTACMTYAACCTGTGHTRTC | CCACCRAAGCAAAAACCAGCT |  | 6270 | 8135 | 1865 | ^b^ |
| 9 | GGTAAGTGCATGACTTTYGATGC | CCACCRAAGCAAAAACCAGCT |  | 7191 | 8135 | 944 |  |
| 9 to 11 | GGTAAGTGCATGACTTTYGATGC | ACCYACTGACCACARGTACCAGC |  | 7191 | 9342 | 2151 |  |
| 10 to 14 | TGGTTATTAAGAAYGGTRTYGTTCAACC | TGYCTAAYCTTGGAAAGCACTTCA |  | 7567 | 11363 | 3796 |  |
| 14 to 16 | GCTTACCATGTTGATAAGTCTTACTACAAA | AGACTTGCTCATGGTCCATAACG |  | 10313 | 12501 | 2188 |  |
| 17 | TTAAACGAGTGCGGGGTTCTAG | GGYACACCATCWATRTGRACTTTACG |  | 12282 | 13264 | 982 |  |
| 18 to 19 | ATGGGAATKACTTCATGTTTAGAR | CAGAAACAGCTTGAAAGATGT |  | 12977 | 14352 | 1375 |  |
| 20 | ACAACYAGYGGTGATGGTACTACA | CCAAACACATTACCATTAGCACAGAG |  | 14285 | 15316 | 1031 |  |
| 21 to 22 | GGCATGTGTGTTGTTTGTGGTT | TGTARATDACATAATCRTACTCACTACC |  | 15050 | 16684 | 1634 |  |
| 23 | TGCCCAGCTGARATTGTTAARACAG | CATAGTTGTAAGYTCAAGACCACC | ORF1b region | 16376 | 17426 | 1050 |  |
| 24 to 25 | TTTGCTATGCGTAATGTDAGAGCRTG | TGGCACTRAGYCCYTTCACAGC |  | 17072 | 18835 | 1763 |  |
| 26 to 27 | AATRGYAAAGCMCTCCARAGT* | GARTTTCKCCAAAATATATAATTGGCATGC |  | Not present | 20120 | 1648 | ^c^ |
| 28 to 32 | GACGCAGACTTCAGTGTTA | ACCATTATGCCATTRTARTA | Spike | 19728 | 23341 | 3613 | ^d^ |
| 33 | GCCCTTAAYCTTGGYGCRCGTMT | ACRCATATTCCTGACCATGCCG |  | Not present | Not present |  | ^e^ |
| 34 to 35 | GCAGCACTTAAYGCBTATGYGTC | AGCATAGGGTCTACAAAATGCAAC |  | Not present | 24616 |  | ^f^ |
| 35 | TGTCTBAGTACTGGHTGYTGTGG | AGCATAGGGTCTACAAAATGCAAC | ORF3c/E/M region part 1 | 23592 | 24616 | 1024 |  |
| 36 | GATGGCATTGTKACAAYAACTGTCTT | ARCCGAACATTACATATCTGGAAACTT | ORF3c/E/M region part 2 | 24380 | 25489 | 1109 |  |
| 36 to 37 | GATGGCATTGTKACAAYAACTGTCTT | CTTATTACCTATTCCYTTGGGAACAA |  | 24380 | 25407 | 1027 |  |
| 38 | CYGGTGATTACTCAACAGAAGCA | GTTTTGGCATCATCYTTGGCAGG |  | 25779 | 26805 | 1026 |  |
| 38 to 40 | CYGGTGATTACTCAACAGAAGCA | ACATTTTAAACAATCACTAGATCCAGACG |  | 25779 | 28167 | 2388 |  |
| 40 | CARCTTTTGARRCCAGACTGYC | ACATTTTAAACAATCACTAGATCCAGACG |  | 27161 | 28167 | 1006 |  |

^a^Also amplifies GHR, deletion variant; shorter. ^b^Poor performance. ^c^Amplification works, however F primer was not incorporated into the draft genome possibly due to variation in the amplified viruses. ^d^ Also amplifies GHR, deletion variant shorter. ^e^ Poor performance. ^f^ Very poor performance

**Supplementary table S20: Tiled amplicon scheme for the amplification of the FCoV-23 genome.** Primers were designed using primal scheme^1^ and manual design using a composite FCoV-23 genome generated using primers in supplementary table 20. Amplicon lengths are designed for 800 bp lengths with 80 bp overlap. We recommend to first assess presence of full-length (FL) spike or domain 0 deletion using primers 36A-F and

| Primer number | Forward (5'-3') | Location (FL genome) | LONG Pool | SHORT Pool | x-fold in Pool |
| --- | --- | --- | --- | --- | --- |
| 1-F | GGACACCAACTCGAACTAAACGA |  | 1 | 1 | 0.5 |
| 1-R | ARTTCTTCTTGACTGGTGGTGC |  | 1 | 1 | 0.5 |
| 2-F | ACGGARTAAGTGATCTTAAACCTGTTCT |  | 2 | 2 | 0.5 |
| 2-R | GAAGTCGTTCAATTCGCCTTCAAA |  | 2 | 2 | 0.5 |
| 3-F | TCTTTGCTAAYAGTGTGCTCCAA |  | 1 | 1 | 1 |
| 3-R | AAGCCATATCACCAAYAAYGACA |  | 1 | 1 | 1 |
| 4-F | GCRCTTGTYAAGCTTGTCAGTG |  | 2 | 2 | 1 |
| 4A-F | TGACACCACAGAGGACTGAGGC |  | 2 | 2 | 1 |
| 4-R | AACYTTAAYACCATTACCTAGC |  | 2 | 2 | 1 |
| 4A-R | TCCARTTCCAGCAGAAGGTCWG |  | 2 | 2 | 1 |
| 5-F | CCAGTKTGTCTTAAAAACCATGTYGG |  | 1 | 1 | 1 |
| 5-R | GTACCAATGACYTTTTCAAGCACC |  | 1 | 1 | 1 |
| 6-F | AGAGATASAACCYGTTACACGTGTC |  | 2 | 2 | 1 |
| 6-R | TCGTCTCCAAAAGRTATTCTGCATC |  | 2 | 2 | 1 |
| 7-F | GATGAACAGGAAKCTGAACAACC |  | 1 | 1 | 1 |
| 7A-F | GAYCCATGGGCTGCTGCTGTT |  | 1 | 1 | 1 |
| 7-R | TCCKTGGTARAATGARACTTTTCCC |  | 1 | 1 | 1 |
| 7A-R | CTTTGACGTTCTTCTGCTACRCAC |  | 1 | 1 | 1 |
| 8-F | TGGATGGTATGGGAATTAAACCTCG |  | 2 | 2 | 1 |
| 8-R | CACACTWGGYAACTGGTTAGT |  | 2 | 2 | 1 |
| 9-F | TGTCTTYGTTTACACTGACCARGAG |  | 1 | 1 | 1 |
| 9-R | CGTARTAGGTGTAATGACCACGYG |  | 1 | 1 | 1 |
| 10-F | GCTTGKTGAKTTGATGTCRAGTG |  | 2 | 2 | 2 |
| 10-R | ACTAGTTTTGCATARCGCCA |  | 2 | 2 | 2 |
| 11-F | GCTGACGTRTTCTTTATGRCTGG |  | 1 | 1 | 1 |
| 11-R | GCTTTTTRCAGAGTYTACCAGTACC |  | 1 | 1 | 1 |
| 12-F | TCATGGGATTAYAAGTCAGACCC |  | 2 | 2 | 1 |
| 12-R | TTGGCATTRAARAGTGCTCCCT |  | 2 | 2 | 1 |
| 13-F | CARGAAGTGCTTAAGACTATGTTYC |  | 1 | 1 | 1 |
| 13A-F | TTCACGGCATATGACTATGATG |  | 1 | 1 | 1 |
| 13B-F | GAAGTATAGTAGTCAGGAAGTGC |  | 1 | 1 | 2 |
| 13-R | CCATGTAATCATARCCCTCWGCAG |  | 1 | 1 | 2 |
| 13A-R | CAAATGGTTGAACAACACCGTTC |  | 1 | 1 | 2 |
| 14-F | CAGATGAAGATYTGCCKTATGAGCG |  | 2 | 2 | 2 |
| 14-R | CCATCYCCAAACTCTTTATCATAGACA |  | 2 | 2 | 2 |
| 14A-R | TCAATACACTCTCCGACTCTGC |  | 2 | 2 | 2 |
| 15-F | TGAGGGTGCTAAGCTTTACAGTG |  | 1 | 1 | 0.5 |
| 15-R | ATCAGCCTCTCCCATMGAACCA |  | 1 | 1 | 0.5 |
| 16-F | TCYCTACCATCACTATTCAAACTTAARGT |  | 2 | 2 | 1 |
| 16-R | GAGAGCCGTTACCTAATTCTAGATGG |  | 2 | 2 | 1 |
| 17-F | GTCTGTGAAACCAGGTGAGAGTT |  | 1 | 1 | 0.5 |
| 17-R | AACAACATTTTATGCTTAATTCCAACGAC |  | 1 | 1 | 0.5 |
| 18-F | ACGCCTACTGAAGTCATAAGGC |  | 2 | 2 | 0.5 |
| 18-R | ACTGTGAACTGGTAAACACCACA |  | 2 | 2 | 0.5 |
| 19-F | CAAAGGACTGGTTTGTTGTTTTTGC |  | 1 | 1 | 0.5 |
| 19-R | GCGTGCCTCTTTGTACATGCYA |  | 1 | 1 | 0.5 |
| 20-F | TTGCCTAGCTGGATTGCCTATG |  | 2 | 2 | 1 |
| 20-R | TATTTAACYTCAGGACCATTARCRCC |  | 2 | 2 | 1 |
| 21-F | TGCTTATGGTAGYGGTAAAGCGC |  | 1 | 1 | 0.5 |
| 21-R | AGCTCTACTAACATGGTCTGGATC |  | 1 | 1 | 0.5 |
| 22-F | GWGGTATGCAGCCAGTTAMTAAYT |  | 2 | 2 | 1 |
| 22-R | ACTGGRTCAAACCAATCCTTRTT |  | 2 | 2 | 1 |
| 23-F | TGYAGTTGCTGAACAYGACTT |  | 1 | 1 | 1 |
| 23-R | GKAGATCTGTCATRGTCAACTTCA |  | 1 | 1 | 1 |
| 24-F | CATTGTGCYAATTTTAACACRYT |  | 2 | 2 | 0.5 |
| 24A-F | CTGCATTTGGACCTCTTGTACGTA |  | 2 | 2 | 0.5 |
| 24-R | CTGTCTCGTYGTCATTGTKGA |  | 2 | 2 | 0.5 |
| 24A-R | TCCCACAGTGCGAGCTCTAGAC |  | 2 | 2 | 0.5 |
| 25-F | TTTGGTAAGGCAAGACTTTACTATGAGA |  | 1 | 1 | 1 |
| 25-R | TCTGCTACATARCCAAGATCWGCA |  | 1 | 1 | 1 |
| 26-F | GCTTTTAGGAGTRGATTCAAACAC |  | 2 | 2 | 1 |
| 26A-F | ACAGCGCAAGATATATGACAATTG |  | 2 | 2 | 1 |
| 26-R | AGCTACACACATAWGGYGTAATAGAC |  | 2 | 2 | 1 |
| 26A-R | GTAACATCATTAACAGTACAACCATT |  | 2 | 2 | 1 |
| 27-F | CAAAACACCCTAARCCTGCWTATCAA |  | 1 | 1 | 0.5 |
| 27-R | GAGTCACTACCATATTCTGACTGCTC |  | 1 | 1 | 0.5 |
| 28-F | TGTGAAAGCWAAGGAGGARTCTGT |  | 2 | 2 | 1 |
| 28-R | AARGTTCTAGGWGCTGGRAGTT |  | 2 | 2 | 1 |
| 29-F | AGGATAATACCTCAAAGAATCAGAGTTGA |  | 1 | 1 | 1 |
| 29-R | ACCACAAGTTTCAGGTTTTGCYTG |  | 1 | 1 | 1 |
| 30-F | ACTCGGCKCAAGGTAGTGAGTA |  | 2 | 2 | 0.5 |
| 30-R | CCATTTTGCCGCAMTCACATTT |  | 2 | 2 | 0.5 |
| 31-F | TCATGAGGAGRGGTCAAYCYT |  | 1 | 1 | 1 |
| 31-R | GCATGATTGTTAACATACAACGCAC |  | 1 | 1 | 1 |
| 32-F | AAYAATGTTAGATGTCTGGAGTAYGA |  | 2 | 2 | 1 |
| 32-R | AGTTTGAGAAAGGACAGTCCGC |  | 2 | 2 | 1 |
| 33-F | AGGAACGGACCTACTGACAAGT |  | 1 | 1 | 0.5 |
| 33-R | TCGTCCAAGAGTATGTCCATATAAGTG |  | 1 | 1 | 0.5 |
| 34-F | TGGTTTTGAACACGTTGTATTTGGA |  | 2 | 2 | 0.5 |
| 34-R | ATCACCTGTAACACTGAAGTCTGC |  | 2 | 2 | 0.5 |
| 35-F | GCTCCTGGTAGTACTGTCYTAAGA |  | 1 | 1 | 1 |
| 35A-F | GTCAAGTACACTCAGTTGTGTC |  | 1 | - | 1 |
| 35-R | CACCAACAACYACASTTCCTTCT |  | 1 | - | 1 |
| 35A-R | CGTTGCCATCTAATTGTGTTACG |  | 1 | - | 1 |
| 36-F | GGCAARYTACTAAACTTTGGTAACC |  | 2 | 2 | 1 |
| 36A-F | CTAAGGAAGGGTAAGTTGCTCA |  | 2 | 2 | 1 |
| 36-R | CCAGTGCAATRTTCATAATCYTCACA |  | 2 | 1 | 1 |
| 36A-R | CCATCAGGTATGTAACCTCCC |  | 2 | 1 | 1 |
| 37-F | GTGGAATGATGAMYTTGTTACAGC |  | 1 | - | 1 |
| 37A-F | ATACCCACGGACAATGGAACGA |  | 1 | - | 1 |
| 37-R | AGGAAATGTGCTAAAGAAATTGTAACCA |  | 1 | 2 | 1 |
| 37A-R | CCTTTACACTAGGTGGTAATGTTC |  | 1 | 2 | 1 |
| 38-F | ACAGTGMGTGAGTCTAGTTYTTACA |  | 2 | 1 | 1 |
| 38-R | CARTTAGCACCAACAGGAYKCA |  | 2 | 1 | 1 |
| 39-F | ACGTGTATTGCATTCGTTCTAATCAA |  | 1 | 2 | 0.5 |
| 39-R | GTATCGTGACATTACCRGTGCT |  | 1 | 2 | 0.5 |
| 40-F | GTRACRCCATGTGATGTAAGCGC |  | 2 | 1 | 2 |
| 40-R | TGCCATTGTAATATTGAGCACAMAC |  | 2 | 1 | 2 |
| 41-F | GTGACATCTGGYTTAGGTACAGTCG |  | 1 | 2 | 1 |
| 41-R | AACRAGTCCRAAAGTGYGATCG |  | 1 | 2 | 1 |
| 42-F | ACWAGCAGAGGTTAGGGCTAGT |  | 2 | 1 | 0.5 |
| 42-R | TCTACTGAAAAGAGAATGACAACAGCT |  | 2 | 1 | 0.5 |
| 43-F | GTCTYAGTACTGGYTGTTGTGG |  | 1 | 2 | 1 |
| 43-R | GTTRTTGTMACAATGCCATCTATGT |  | 1 | 2 | 1 |
| 44-F | ACCACATGTTAATACCATAGTACAACAAC |  | 2 | 1 | 0.5 |
| 44-R | CATGGCGTGCAGGTAGTACTAT |  | 2 | 1 | 0.5 |
| 45-F | CGTGTCTATGATGTTTCCTAGGGC |  | 1 | 2 | 0.5 |
| 45-R | TGTAGGTRTGCCATCAAGGGGT |  | 1 | 2 | 0.5 |
| 46-F | TCGGCTTTAGTGTTGCAGGTG |  | 2 | 1 | 0.5 |
| 46-R | AGTAGAAGAACCACCTTTCAGGAAG |  | 2 | 1 | 0.5 |
| 47-F | CCCATTACCCTCGAAACAGGATC |  | 1 | 2 | 0.5 |
| 47-R | GTRAGYGTGACTTTCACYTGATC |  | 1 | 2 | 0.5 |
| 48-F | ATGCCAACAAACACASCTGG |  | 2 | 1 | 0.5 |
| 48-R | CACACWAGGAYTACARCAATCATG |  | 2 | 1 | 0.5 |
| 49-F | CCTGCTATACATTGTTAGGTGC |  | 1 | 2 | 1 |
| 49-R | ACATTTTAAACAATCACTAGATCCAGACG |  | 1 | 2 | 1 |

**Supplementary table S21: Results of recombination analysis.** Several tools run using RDP5 and using a multi sequence alignment between the assembled FCoV-23 genome 2-C11 Re 10276 (GB reposited – no number assigned yet), a pCCoVII genome (KP981644.1), an FCoVII genome (LC742526.1) and an FCoVI genome (MT239440.1) were used to analyse recombination.

| Tool used via RDP5 | P- values |
| --- | --- |
| RDP5^4^ | 4e-30 x10^-300^ |
| GENECONV | 4e-30 x10^-300^ |
| BootScan^5^ | 8.624 x10^-269^ |
| MaxChi^6^ | 1.200 x10^-199^ |
| Chimaera^7^ | 2.948 x10^-64^ |
| SiScan^8^ | 8.209 x10^-93^ |
| 3Seq^9^ | 4e-30 x10^-300^ |

**Supplementary table S22: Results of recombination analysis.** Several tools run using RDP5 and using a multi sequence alignment between the assembled domain 0-deletion FCoV-23 genome 2-F12 BW 11350 (GB reposited – no number assigned yet), a pCCoVII genome (KP981644.1), an FCoVII genome (LC742526.1) and an FCoVI genome (MT239440.1) were used to analyse recombination.

| Tool used via RDP5 | P- values |
| --- | --- |
| RDP5^4^ | 2.000 x10^-299^ |
| GENECONV | 2.000 x10^-299^ |
| BootScan^5^ | 4.414 x10^-21^ |
| MaxChi^6^ | 6.000 x10^-199^ |
| Chimaera^7^ | 6.000 x10^-199^ |
| SiScan^8^ | 9.679 x10^-87^ |
| 3Seq^9^ | 2.000 x10^-299^ |

**Supplementary table S23: Comparison of FCoV-23 Spike-2 with determinant mutations.** The consensus FCoV-23 sequence was compared with key sequence features that were identified in Zehr *et al.* ^10^ as positively associated with the FECV biotype. Association with biotype was assessed whether mutations had been previously more likely been associated with one of the biotypes. Amino acid positions below the line are after the recombination breakpoint between pCCoV and FCoV-1.

| Position in FCoV-23 | Amino acid sequence in FCoV-23 | Amino acid composition at site associated with biotype ^10^ |
| --- | --- | --- |
| 534 | V | Marginally tentative FIPV |
| 596 | Q | Marginally tentative FIPV |
| 1404 | L | **New mutation** |
| 1405 | V | Marginally tentative FIPV |
| 1416 | L | Marginally tentative FECV |
| 1434 | L | Marginally tentative FECV |

**Supplementary table S24: Comparison of FCoV-23 Orf3a,b, and c with determinant mutations.** The consensus FCoV-23 sequence was compared with key sequence features that were identified in Zehr *et al.* ^10^ as positively associated with the FECV biotype. Association with biotype was assessed whether mutations had been previously more likely been associated with one of the biotypes. Sites could previously not be “statistically associated uniquely with one phenotype”^10^.

| Protein | Position in FCoV-23 | Amino acid sequence in FCoV-23 | Amino acid composition at site associated with biotype ^10^ |
| --- | --- | --- | --- |
| Orf3a | **30** | L | No indication |
| Orf3a | **32** | N | No indication |
| Orf3a | **47** | E | No indication |
| Orf3a | **58** | Q | No indication |
| Orf3a | **61-62 gap** | IE | No indication |
| Orf3a | **64** | S | No indication |
| Orf3a | **65** | S | **New mutation** |
| Orf3b | **2** | R | **New mutation** |
| Orf3b | **64** | K | No indication |
| Orf3b | **71** | A | No indication |
| Orf3c | **11** | S | No indication |
| Orf3c | **71** | G | No indication |
| Orf3c | **72** | V | No indication |
| Orf3c | **159** | M | No indication |
| Orf3c | **165** | T | No indication |
| Orf3c | **175** | G | No indication |

**Supplementary table S25: Comparison of FCoV-23 Orf7b with determinant mutations.** The consensus FCoV-23 sequence was compared with key sequence features that were identified in Zehr *et al.* ^10^ as positively associated with the FECV biotype. Association with biotype was assessed whether mutations had been previously more likely been associated with one of the biotypes.

| Position in FCoV-23 | Amino acid sequence in FCoV-23 | Amino acid composition at site associated with biotype ^10^ |
| --- | --- | --- |
| 5 | V | Marginally tentative FIPV |
| 11 | L | No indication |
| 12 | A | No indication |
| 19 | D | No indication |
| 25 | H | No indication |
| 36 | Q | No indication |
| 39 | V | No indication |
| 41 | H | No indication |
| 48 | H | No indication |
| 50 | I | No indication |
| 63 | S | No indication |
| 68 | N | No indication |
| 82 | I | No indication |
| 89 | S | No indication |
| 106 | N | No indication |
| 107 | Q | No indication |
| 129 | T | No indication |
| 131 | F | No indication |
| 139 | T | No indication |
| 140 | Q | No indication |
| 145 | R | No indication |
| 147 | F | No indication |
| 149 | H | No indication |
| 152 | S | No indication |
| 159 | **I** | **New mutation** |
| 160 | H | No indication |
| 167 | Y | No indication |
| 168 | C | No indication |
| 170 | H | No indication |
| 172 | L | No indication |
| 187 | K | No indication |
| 190 | R | No indication |
| 191 | S | No indication |
| 194 | V | No indication |
| 198 | L | No indication |
| 199 | N | No indication |
| 200 | Q | No indication |
| 202 | H | No indication |
| 203 | **H** | **New mutation** |
| 204 | T | No indication |
